# *De novo* coding variants in the *AGO1* gene cause a neurodevelopmental disorder with intellectual disability

**DOI:** 10.1101/2020.12.23.424103

**Authors:** Audrey Schalk, Margot A. Cousin, Thomas D. Challman, Karen E. Wain, Zöe Powis, Kelly Minks, Aurélien Trimouille, Eulalie Lasseaux, Didier Lacombre, Chloé Angelini, Vincent Michaud, Julien Van-Gils, Nino Spataro, Anna Ruiz, Elizabeth Gabau, Elliot Stolerman, Camerun Washington, Raymond J. Louie, Brendan C Lanpher, Jennifer L. Kemppainen, A. Micheil Innes, R. Frank Kooy, Marije Meuwissen, Alice Goldenberg, François Lecoquierre, Gabriella Vera, Karin E M Diderich, Beth Rosen Sheidley, Christelle Moufawad El Achkar, Meredith Park, Fadi F. Hamdan, Jacques L. Michaud, Ann J. Lewis, Christiane Zweier, André Reis, Matias Wagner, Heike Weigand, Hubert Journel, Boris Keren, Sandrine Passemard, Cyril Mignot, Koen L.I. van Gassen, Eva H. Brilstra, Gina Itzikowitz, Emily O’Heir, Jake Allen, Kirsten A. Donald, Bruce R. Korf, Tammi Skelton, Michelle L Thompson, Nathaniel H. Robin, Natasha Rudy, William B. Dobyns, Kimberly Foss, Yuri A Zarate, Katherine A. Bosanko, Yves Alembik, Benjamin Durand, Frédéric Tran Mau-Them, Emmanuelle Ranza, Xavier Blanc, Stylianos E. Antonarakis, Kirsty McWalter, Erin Torti, Francisca Millan, Amy Dameron, Mari J. Tokita, Michael T. Zimmermann, Nikita R. Dsouza, Eric W. Klee, Amélie Piton, Bénédicte Gerard

## Abstract

High-impact pathogenic variants in more than 1,000 protein-coding genes cause Mendelian forms of neurodevelopmental disorders (NDD), including the newly reported *AGO2* gene. This study describes the molecular and clinical characterization of 28 probands with NDD harboring heterozygous *AGO1* coding variants. *De novo* status was always confirmed when parents were available (26/28). A total of 15 unique variants leading to amino acid changes or deletions were identified: 12 missense variants, two in-frame deletions of one codon, and one canonical splice variant leading to a deletion of two amino acid residues. Some variants were recurrently identified in several unrelated individuals: p.(Phe180del), p.(Leu190Pro), p.(Leu190Arg), p.(Gly199Ser), p.(Val254Ile) and p.(Glu376del). *AGO1* encodes the Argonaute 1 protein, which functions in gene-silencing pathways mediated by small non-coding RNAs. Three-dimensional protein structure predictions suggest that these variants might alter the flexibility of the AGO1 linkers domains, which likely would impair its function in mRNA processing. Affected individuals present with intellectual disability of varying severity, as well as speech and motor delay, autistic behavior and additional behavioral manifestations. Our study establishes that *de novo* coding variants in *AGO1* are involved in a novel monogenic form of NDD, highly similar to *AGO2* phenotype.

## INTRODUCTION

Neurodevelopmental disorders (NDD), such as intellectual disability (ID) and autism spectrum disorder (ASD), have important genetic contributions characterized by extreme heterogeneity. More than a thousand genes have now been implicated in monogenic forms of NDD^1^. Also, many more genes have been identified as candidates for ID/ASD, including genes showing enrichment of rare *de novo* variants in large-scale sequencing studies performed in affected individuals^2^. In individuals affected by ID or ASD, few *de novo* missense variants have been reported in *AGO1* (or *EIF2C1*), a gene encoding a protein from the argonaute family, which participate in RNA silencing pathways, suggesting that *AGO1* could be a promising candidate gene for NDD^3–6^.

The argonaute protein family, identified originally in plants, includes AGO and PIWI proteins, and is involved in gene-silencing pathways guided by small non-coding RNAs (sncRNA, including short interfering RNAs, siRNAs; microRNAs, miRNAs; Piwi-interacting RNAs, piRNAs)^7^. PIWI proteins are involved in transposon repression in germinal cells whereas AGO proteins are involved in translation repression and degradation of targeted mRNA^8^. In addition to their role in mRNA post-transcriptional regulation in the cytoplasm, AGO proteins have also been shown to have nuclear activities, playing a role in the regulation of transcription, chromatin remodeling, alternative splicing regulation, and even in DNA double-strand break repair^8–12^.

Large deletions of 1.1 Mb to 3.1 Mb at the 1p34.3 loci including *AGO1* together with *AGO3* (and sometimes *AGO4* among other genes) were previously reported in children with NDD^13,14^. These five individuals presented with psychomotor developmental delay as well as additional non-specific features (feeding difficulty, language impairment, facial dysmorphy). In addition, *AGO2* was very recently implicated in NDD^15^: 21 individuals with heterozygous *de novo* mutations in *AGO2* variant were reported, including 11 missense variants, one in-frame deletion and one 235 kb deletion involving the first three exons of *AGO2*. Functional studies revealed that those variants hampered correct sncRNA-mediated silencing, having a loss of function effect.

We here report 28 individuals from 26 families affected by NDD carrying heterozygous amino acid substitutions or deletions predicted-damaging in *AGO1*, thus, causally-linking coding variants in this gene to ID. These variants distributed along the different domains of the protein, are predicted to affect the flexibility of AGO1 structure, which might impair its role in sncRNA-induced gene regulation. Several *AGO1* variants reported in this study affect homologous residues to those recently described in *AGO2* and faithfully recapitulate the functional prediction effects. The high similarity in clinical presentation reinforces the description of *AGO1/AGO2* associated phenotype.

## MATERIALS AND METHODS

### Cohort

All affected individuals were recruited independently through a worldwide collaborative network of clinical and molecular geneticists connected notably via GeneMatcher^16^. Affected individuals were referred by clinical genetics services from across Belgium (family 8), Canada (families 7, 12), France (families 3, 9, 16, 17, 20 and 24), Germany (families 14 and 15), Switzerland (family 26), USA (families 1, 2, 5, 6, 11, 13, 19, 21, 22, 23 and 25), Spain (family 4) and The Netherlands (families 10 and 18). Blood samples were obtained following the provision of informed consent by the individuals or their legal representatives. Ethical approval was obtained from the local ethics committees.

### Identification of AGO1 variants

*AGO1* variants were identified by laboratories using various high throughput sequencing strategies (targeted genes panel, exome sequencing or whole genome sequencing) (see **Table 1**). Reads alignment was performed against the human genome reference build GRCh37/hg19; Filtering strategies and description of bioinformatics tools are presented in supplementary data (**Supplementary text**). *AGO1* variants were annotated following the NM_012199.4. Variants were confirmed by independent Sanger sequencing and concordance of the trio (father, mother, child) was confirmed for all *de novo* variants, either by verifying rare SNP inheritance pattern in the case of NGS trios or by microsatellites segregation study, except for families 23 and 25, as no parental samples were available. Predictions of missense variant effects were performed using *in silico* tools such as Combined Annotation Dependent Depletion (CADD^17^), SIFT^18^ and Polyphen-2^19^. Effects on splicing were analyzed using SpliceAI^20^, NNsplice^21^ and MaxEnt^22^ and confirmed, if needed, by cDNA analysis from blood mRNA.

**Table 1:** List of missense changes identified in AGO1 in individuals with intellectual disability. Version of the programs used: SIFT version v6.2.0; Polyphen2 (PPH2) HumVar version v2.2.2r398; CADD version GRCh37-v1.4; †Change in folding free energy upon each mutation, using a 3D structure-based algorithm. See Methods for details. ‡3D molecular context of the amino acid position. See Results for details. αMD simulations were summarized using PC analysis; we summarized the change in PC1 and PC2 using the number of standard deviations away from the average WT conformation that each mutation sampled on average.

| Family | Family previously reported | Genomic position (GRCh37) - chr 1 | cDNA | Protein | SIFT | PPH2 (HumVar) | CADD | Domain | $\Delta\Delta G_{\text{fold}}^{\dagger}$ | Context<sup>‡</sup> | Dynamics Change<sup>α</sup> |
| --- | --- | --- | --- | --- | --- | --- | --- | --- | --- | --- | --- |
| 1, 2, 3, 4, 5 | - | g.36359301_36359303del | c.539_541del | p.(Phe180del) | NA | NA | NA | L1 linker | NA | Core | NA |
| 6 | - | g.36359328C>T | c.566C>T | p.(Pro189Leu) | Deleterious (score: 0.02) | Possibly Damaging (score: 0.605) | 28.3 | L1 linker | 1.0±0.4 | PAZ | PC1 -0.4, PC2 -1.6 |
| 7, 8 | - | g.36359331T>G | c.569T>G | p.(Leu190Arg) | Deleterious (score: 0) | Probably Damaging (score: 1.000) | 29.2 | L1 linker | 9.5±0.6 | PAZ | NA |
| 9 | Rauch *et al.*<sup>3</sup> | g.36359331T>C | c.569T>C | p.(Leu190Pro) | Deleterious (score: 0) | Probably Damaging (score: 1.000) | 29.3 | L1 linker | 5.6±0.4 | PAZ | PC1 -0.3, PC2 -1.1 |
| - | Martinez *et al.*<sup>23</sup> | g.36359345G>A | c.583G>A | p.(Glu195Lys) | Deleterious (score: 0) | Probably Damaging (score: 0.999) | 30 | L1 linker | 2.5±0.3 | PAZ | PC1 0.1, PC2 -0.7 |
| 10, 11, 12 (Hamdan *et al.*)<sup>5</sup>, 13, 14 | Sakaguchi *et al.*<sup>6</sup> | g.36359357G>A | c.595G>A | p.(Gly199Ser) | Deleterious (score: 0) | Probably Damaging (score: 1.000) | 29.6 | L1 linker | 17.9±0.7 | Near Core | PC1 0.6, PC2 -0.9 |
| - | DDD Study<sup>24</sup> | g.36359409A>T | c.647A>T | p.(Asp216Val) | Deleterious (score: 0) | Probably Damaging (score: 0.996) | 32 | L1 linker | 2.5±0.7 | Bridge R712 | PC1 -0.4, PC2 -0.6 |
| 15 | - | g.36359636A>G | c.650-2A>G (r.650_655del) | p.Val217_Ser218del | NA | NA | NA | L1 linker | NA | Linker and RNA H-bond | NA |
| 16 | - | g.36359746G>A | c.758G>A | p.(Arg253His) | Deleterious (score: 0.03) | Benign (score: 0.173) | 25.6 | PAZ | 0.6±0.6 | Helix Cap | PC1 0.2, PC2 -1.5 |
| 17, 18 | - | g.36359748G>A | c.760G>A | p.(Val254Ile) | Deleterious (score: 0) | Benign (score: 0.092) | 22.9 | PAZ | -1.0±0.0 | Helix Center | PC1 -0.7, PC2 -1.5 |
| 19 | - | g.36360821C>T | c.971C>T | p.(Pro324Leu) | Deleterious (score: 0) | Probably Damaging (score: 1.000) | 31 | PAZ | 2.1±0.1 | PAZ | NA |
| - | Sanders *et al.*<sup>4</sup> | g.36367118C>T | c.1064C>T | p.(Thr355Ile) | Deleterious (score: 0) | Probably Damaging (score: 0.973) | 24.8 | PAZ | 0.4±0.6 | Helix Cap | NA |
| 20 | - | g.36367127A>G | c.1073A>G | p.(Gln358Arg) | Deleterious (score: 0) | Probably Damaging (score: 0.996) | 26.8 | PAZ | 0.3±0.2 | Linker | NA |
| 21, 22 | - | g.36367180_36367182del | c.1121_1123del | p.(Glu376del) | NA | NA | NA | L2 linker | NA | Linker | NA |
| 23 | - | g.36367661A>T | c.1253A>T | p.(Tyr418Phe) | Deleterious (score: 0) | Benign (score: 0.053) | 24.4 | L2 linker | 0.3±0.1 | Linker | PC1 0.7, PC2 -0.9 |
| 24 | - | g.36384011A>T | c.2252A>T | p.(His751Leu) | Deleterious (score: 0) | Probably Damaging (score: 0.982) | 29.1 | PIWI | -1.1±0.8 | RNA H-bond | PC1 0.6, PC2 -1.2 |
| 25 | - | g.36384732C>T | c.2342C>T | p.(Thr781Met) | Deleterious (score: 0) | Probably Damaging (score: 1.000) | 28 | PIWI | 1.8±0.9 | PIWI core | NA |
| 26 | - | g.36384779A>T | c.2389A>T | p.(Ile797Phe) | Deleterious (score: 0) | Probably Damaging (score: 0.969) | 28.3 | PIWI | 3.1±0.5 | PIWI near RNA | PC1 1, PC2 -0.8 |

### Collection of clinical information

Physicians with experience in clinical dysmorphology examined *a posteriori* all 28 individuals of this cohort, including the individual carrying the previously described variant in Hamdan *et al*^5^ (F12). Photographs were collected from research participants after informed consent had been obtained. Clinical data for additional individuals with *de novo* variants reported in previous publications^3–6,23,24^ were retrieved from published data or supplemental data. Face2Gene Research application (FDNA Inc., Boston, MA) using DeepGestalt technology (algorithm 19.1.9)^25^ was used to evaluate the facial characteristics of the individuals with *AGO1* variants. Fourteen frontal facial photos were obtained from unrelated affected individuals (proband family: age at photo: F3: 7 years, F4: 8 years, F5: ∼3-6 years, F6: 12 years, F9 twin 2: 3 years, F10: 8 years, F11: 11 years, F13: 7 years, F14: 5.7 years, F19: 2 years, F20: 3 years, F21: 3.5 years, F23: 9 years, F24 twin1: 10 years) and compared to 14 control images matched for age, sex, and ethnicity provided by Face2Gene. To estimate the power of DeepGestalt in distinguishing affected individuals from controls, a cross validation scheme was used, including a series of binary comparisons between all groups. For these binary comparisons, the data were split randomly multiple times into training sets and test sets. Each set contained half of the samples from the group, and this random process was repeated 10 times. The results of the comparisons are reported using the receiver operating characteristic (ROC) curve and area under the curve (AUC).

### Building 3D Model for AGO1

The protein structure of human AGO1 has been experimentally determined (PDB 4kxt^26^). We used the available experimental structure with homology-based methods^27^ to fill in short sections of the protein that were not resolved in the experiment. Structures were solved with simple (poly A) guide RNAs that were also partially resolved. We added data from the rhotabacter sphaeroides Argonaute experimental structure (PDB 5awh^28^) and used Discovery Studio (BIOVIA. Dassault Systèmes BIOVIA, Discovery Studio Modeling Environment, Release 4.5, San Diego: Dassault Systèmes; 2015) to complete coordinates of a 21-base (A_19_U_2_) guide RNA. We used BioR^29^ to compile protein annotations from multiple sources including dbNSPF^30^. We used FoldX^31^ v4.0 for mutagenesis and structure-based calculations of ΔΔG_fold_. We visualized protein structures using PyMOL (Molecular Graphics System, Version 1.9 Schrödinger, LLC).

### Molecular Dynamics Simulations

Generalized Born implicit solvent molecular dynamics (isMD) simulations were carried out using NAMD^32^ and the CHARMM36 force field, using a similar procedure to our previous work. Briefly, we utilized an interaction cutoff of 12Å with strength tapering (switching) beginning at 10Å, a simulation time step of 1fs, conformations recorded every 2ps. Each initial conformation was used to generate 6 replicates, and each was energy minimized for 5,000 steps, followed by heating to 300K over 300ps via a Langevin thermostat. A further 13ns of simulation trajectory was generated and the final 10ns (60ns per variant) were analyzed.

### Analysis of Protein Structures and Simulations

All trajectories were first aligned to the initial wild type conformation using C^α^ atoms. Root Mean-Square Deviation (RMSD) values were reported for each after aligning to the initial WT conformation. We calculated per-domain RMSDs by first re-aligning all trajectories using each domain individually, then measuring the RMSD within that domain, alone. This was done independently for each domain, providing a measure of distortion within each domain and across MD simulation time. Root Mean-Square Fluctuation (RMSF) values were calculated at the residue level for the whole protein across trajectories aligned to the initial WT conformation. Variances were computed using median absolute difference (MAD). We computed Z-scores of data by MAD-scaling the difference between each observation and the median. We used C^α^ cartesian space Principal Component (PC) analyses across all simulations to define the essential dynamics of AGO1. Data from each variant was projected onto the PCs to compare how essential motion is activated or suppressed by each genomic variant. Individual PCs were visualized using porcupine plots where a cone is placed to represent the direction and relative magnitude of each residue’s motion. We calculated free-energy landscapes (FELs) of MD trajectories using the approach of Karamzadeh, *et al*.^33^. We show topologic lines from the FEL where each line indicates a specific conformational sampling probability and matched by a corresponding color gradient where each color also indicates a specific conformational sampling probability. We calculated internal distances for an alignment-free measure of conformational changes^34^. We used the ensemble-averaged median pairwise distance between residues, calculated on the last half of each trajectory, as a summary of the change in internal distances. To simplify visualization of these median changes, we averaged information across groups of three consecutive amino acids. To conservatively estimate statistical significance, we used a permutation procedure where data were sub-sampled to 100 points and compared using a t-test. This was repeated 1000 times and the median p-value across repeats reported. The analysis was carried out using custom scripts, leveraging VMD and the Bio3D R package. Protein structure visualization was performed in PyMol and VMD.

## RESULTS

### Identification of de novo variants in AGO1 in individuals with intellectual disability

A total of 15 different heterozygous variants in *AGO1*, including 12 amino acid substitutions, two single amino acid deletions, and one splice variant leading to a two-residues deletion were identified in 26 unrelated families (**Table 1, Figure 1**). For all individuals for whom parental DNA was available (all families except 23 and 25), variants were found to occur *de novo*. No additional genetic variant potentially explaining the clinical manifestations was identified in all these individuals (**Supplementary text**). Two of the amino acid changes we identified had already been reported in individuals with NDD: p.(Leu190Pro) (Rauch *et al*.^3^) and p.(Gly199Ser) (Hamdan *et al*.^5^ and Sakaguchi *et al*.^6^). Thirteen variants were novel to this study. The proband from family F15 has a *de novo* c.650-2A>G variant predicted to impair the use of the canonical acceptor splice site (MaxEnt: −100.0%, NnSplice: −100.0%) and to activate a cryptic acceptor site (Splice AI prediction). Activation of the cryptic acceptor site was subsequently confirmed by mRNA Sanger sequencing (**Supplementary Figure S1**), leading to the in-frame deletion of two amino acids (r.650_655del, p.Val217_Ser218). Six variants were recurrent: p.(Gly199Ser) (six individuals), p.(Phe180del) (five individuals), p.(Leu190Arg) (two individuals), p.(Val254Ile)(two individuals) and p.(Glu376del) (two individuals). The remaining variants were unique: p.(Pro189Leu), p.(Arg253His), p.(Val217_Ser218del), p.(Arg253His), p.(Pro324Leu), p.(Gln358Arg), p.(Tyr418Phe), p.(Thr781Mel) and p.(Ile797Phe). Including the three previously reported missense variants: p.(Glu195Lys)^23^, p.(Asp216Val)^24^ and p.(Thr355Ile)^4^, a total of 18 variants were identified in individuals with NDD (**Table 1, Figure 1**). All identified variants were absent from the general population included in the gnomAD database and all but one affect amino acid residues that are highly conserved until *C. elegans* **(Supplementary Figure S1)**. *In silico* predictions using SIFT and Polyphen2 predict all these missense variants to be deleterious except p.(Arg253His) and p.(Val254Ile). A depletion of missense changes is observed in general populations, with three times fewer missense variants observed than expected in gnomAD cohorts (ratio=0.31, Z-score= 5.68). The CADD scores of missense variants identified in individuals with NDD (mean=27.8) are significantly higher than those of missense variants reported in gnomAD (mean: 24.3, t-test p-value=0.0001).

**Figure 1:**
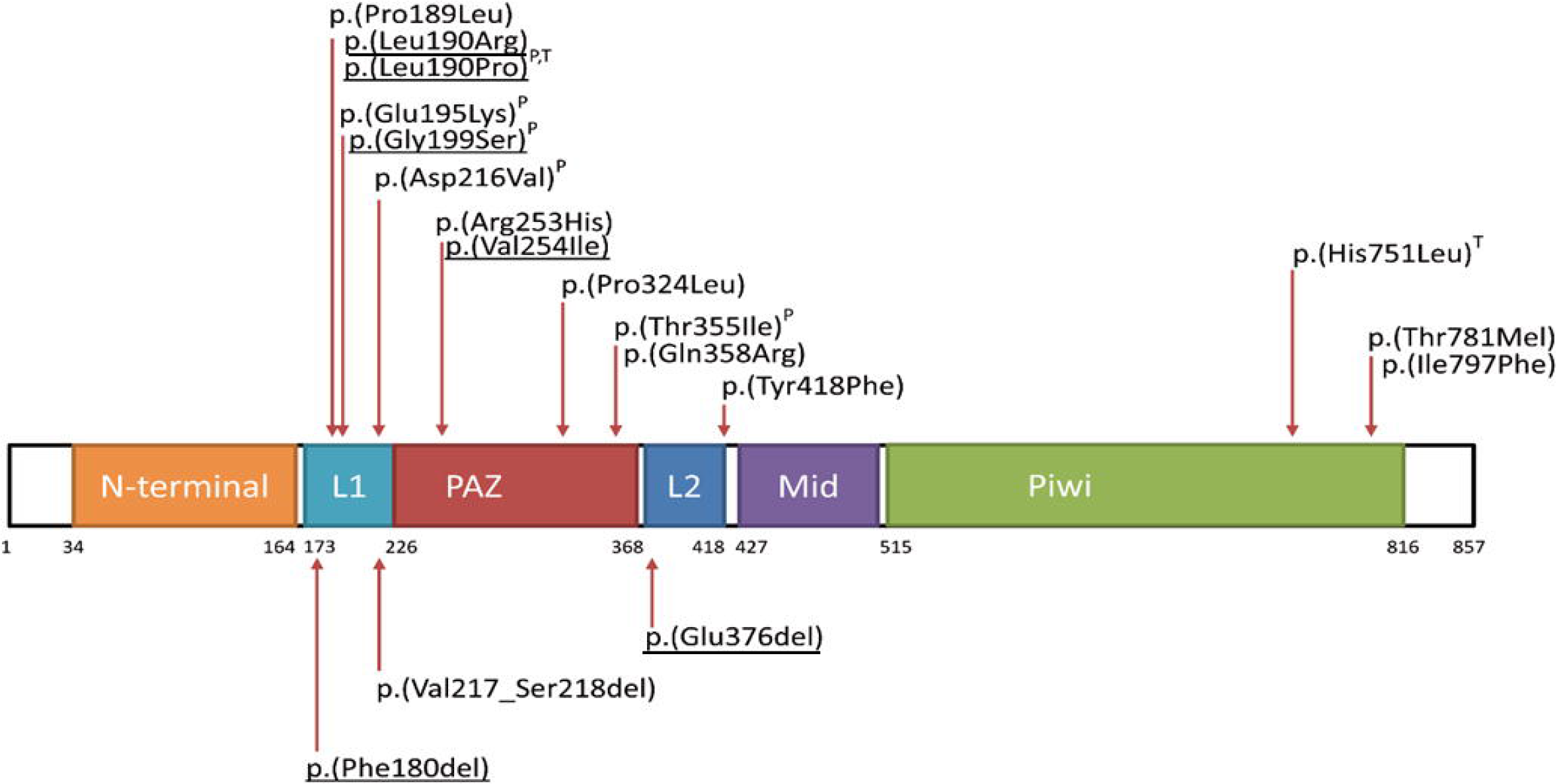
Schematic representation of AGO1 protein with its functional domains showing the locations of the missense variants identified in individuals with ID. AGO1 protein (NP_036331.1) with its functional domains: N-terminal domain (34-164), Linker 1 (L1) domain (173-225), PAZ domain (226-368), Liker 2 (372-418), Mid domain (427-508), and PIWI domain (515-816). PAZ and PIWI domains have motifs of interaction with RNA guide and PIWI domain has motif of RNA blocked access to the active site. The arrows show *de novo* variants identified in individuals from this cohort or previously reported (P: previously reported in literature); T: found in monozygotic twins, and in underlined those that are recurrent.

### Clinical manifestations observed in individuals with AGO1 variants

We collected the clinical characteristics of the 28 affected individuals enrolled in this study: first clinical description for 27 individuals and updated clinical information for one previously reported^5^. We also retrieved the clinical information from previous publications reporting individuals with variants in *AGO1*^3–6,23,24^ and thus obtained clinical data for a total of 33 individuals (17 males, 16 females) from 31 unrelated families. Head circumference and height and weight measurements were in the normal ranges at birth and postnatally, except for three individuals (including two twins) who showed neonatal and postnatal microcephaly (<-2 SD). All affected individuals showed borderline to severe ID, **(Table 2, Supplementary text)** and all showed a severe language delay (30/30; 100 %). Most individuals were able to construct a sentence but with limited spontaneous communication. Three did not acquire language at all and one presented with language regression. A motor developmental delay was also observed in most of the affected individuals: 93 % of individuals had delayed walking (28/30; mean age at onset of independent walking ∼25 months) and seven individuals had persistent hypotonia (13/18; 72 %). Almost half of the individuals had documented seizures or history of seizures (13/28; 46 %), seizures were fever-induced in two families (F7, F18). Twin girls (F9) have different types of seizure: one has photosensitive, drug-resistant seizure and the other has controlled seizure. Interestingly, the three individuals with the most severe epileptic phenotype (as indicated by the history of status epilepticus) share the same variant p.(Gly199Ser) (the fourth individual with this variant has no history of epilepsy at the age of 5 years). Several behavioral features were observed in patients with *AGO1* variants: most of the individuals presented with autistic features (24/30; 80 %), and 11 individuals showed self-harm behavior and/or hetero-aggressiveness (11/14, 78.5%). Attention deficit/hyperactivity was also noticed (15/22, 68 %) and anxiety was reported in seven individuals (7/8, 87.5 %). Moreover, a high percentage of individuals had various sleeping disturbances (17/22, 77 %) including difficulties falling asleep, early awakening, or hypersomnia. Analysis using DeepGestalt technology by Face2Gene did not show significant differences in facial features from matched controls (mean AUC = 0.55 and AUC STD = 0.14, p-value =0.392) **(Figure 2)**. However, if they do not present a recurrent recognizable gestalt, individuals with *AGO1* variants share some subtle common traits such as a thin upper lip, a tall forehead, elongated almond eyes or a small nose. Individuals with feeding difficulties (10/23, 43 %) showed low postnatal weight: a gastrostomy was necessary for six individuals, three individuals had gastro-esophageal reflux and the twins of family 24 received tube feeding during the first three weeks of life in the context of prematurity. Two individuals had recent swallowing problems (F3 and F25). Finally, hypothyroidism was reported in two out of five individuals with the p.(Phe180del) variant. Variable brain MRI anomalies were noticed in 11/24 individuals (46 %), including corpus callosum anomalies (agenesis or hypoplasia), cerebral atrophy, cortical dysplasia or colpocephaly but also non-specific anomalies (increase in extra-axial fluid, T2 and FLAIR signal anomalies, arachnoid cyst).

**Table 2:**
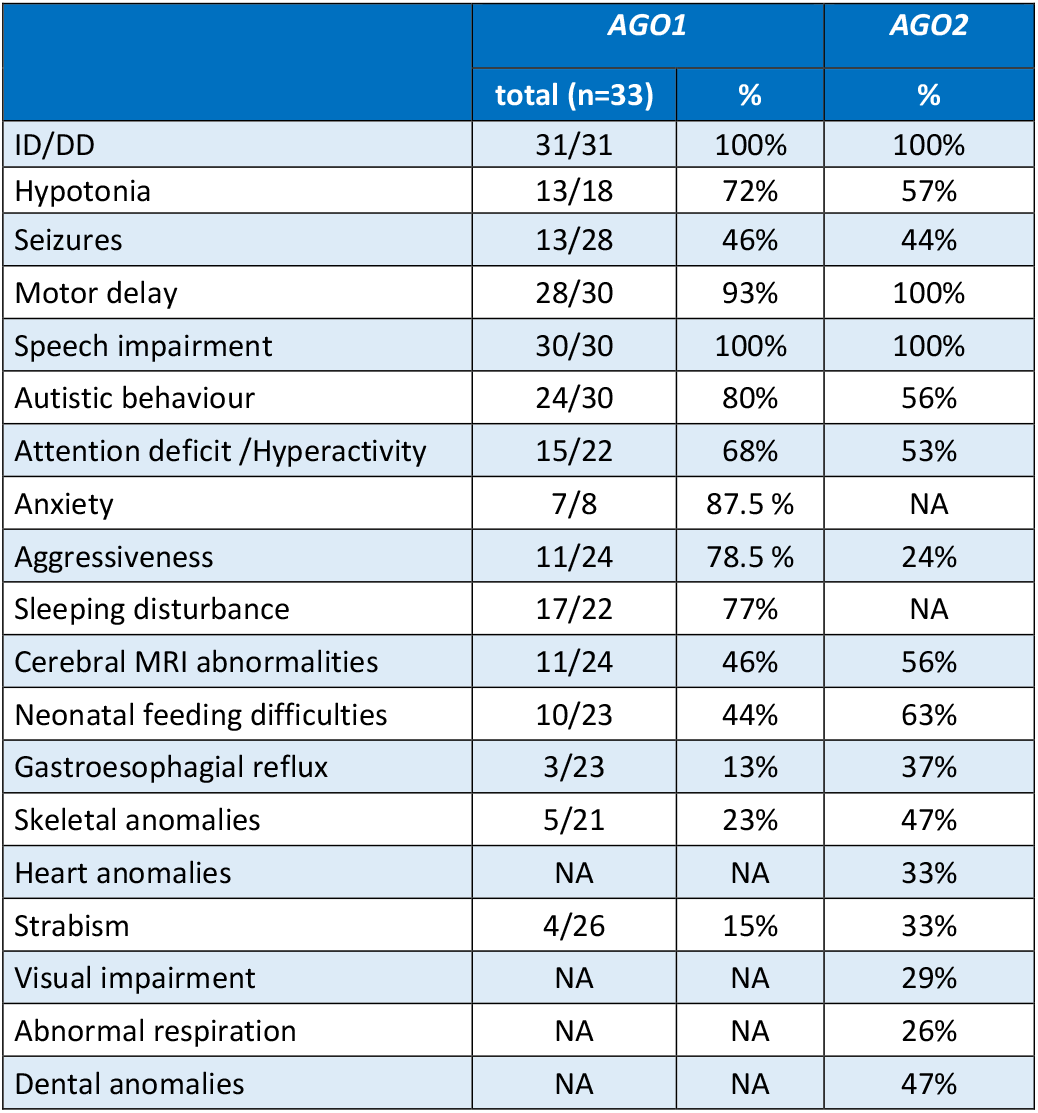
Clinical manifestations observed in individuals with variants in AGO1 (compared to individuals with variants in AGO2 reported by Lessel et al. 2020) ID: intellectual disability; DD: developmental delay; M: male; F: female; NA: not reported

|  | *AGO1* |  | *AGO2* |
| --- | --- | --- | --- |
|  | total (n=33) | % | % |
| ID/DD | 31/31 | 100% | 100% |
| Hypotonia | 13/18 | 72% | 57% |
| Seizures | 13/28 | 46% | 44% |
| Motor delay | 28/30 | 93% | 100% |
| Speech impairment | 30/30 | 100% | 100% |
| Autistic behaviour | 24/30 | 80% | 56% |
| Attention deficit /Hyperactivity | 15/22 | 68% | 53% |
| Anxiety | 7/8 | 87.5 % | NA |
| Aggressiveness | 11/24 | 78.5 % | 24% |
| Sleeping disturbance | 17/22 | 77% | NA |
| Cerebral MRI abnormalities | 11/24 | 46% | 56% |
| Neonatal feeding difficulties | 10/23 | 44% | 63% |
| Gastroesophageal reflux | 3/23 | 13% | 37% |
| Skeletal anomalies | 5/21 | 23% | 47% |
| Heart anomalies | NA | NA | 33% |
| Strabism | 4/26 | 15% | 33% |
| Visual impairment | NA | NA | 29% |
| Abnormal respiration | NA | NA | 26% |
| Dental anomalies | NA | NA | 47% |

**Figure 2:**
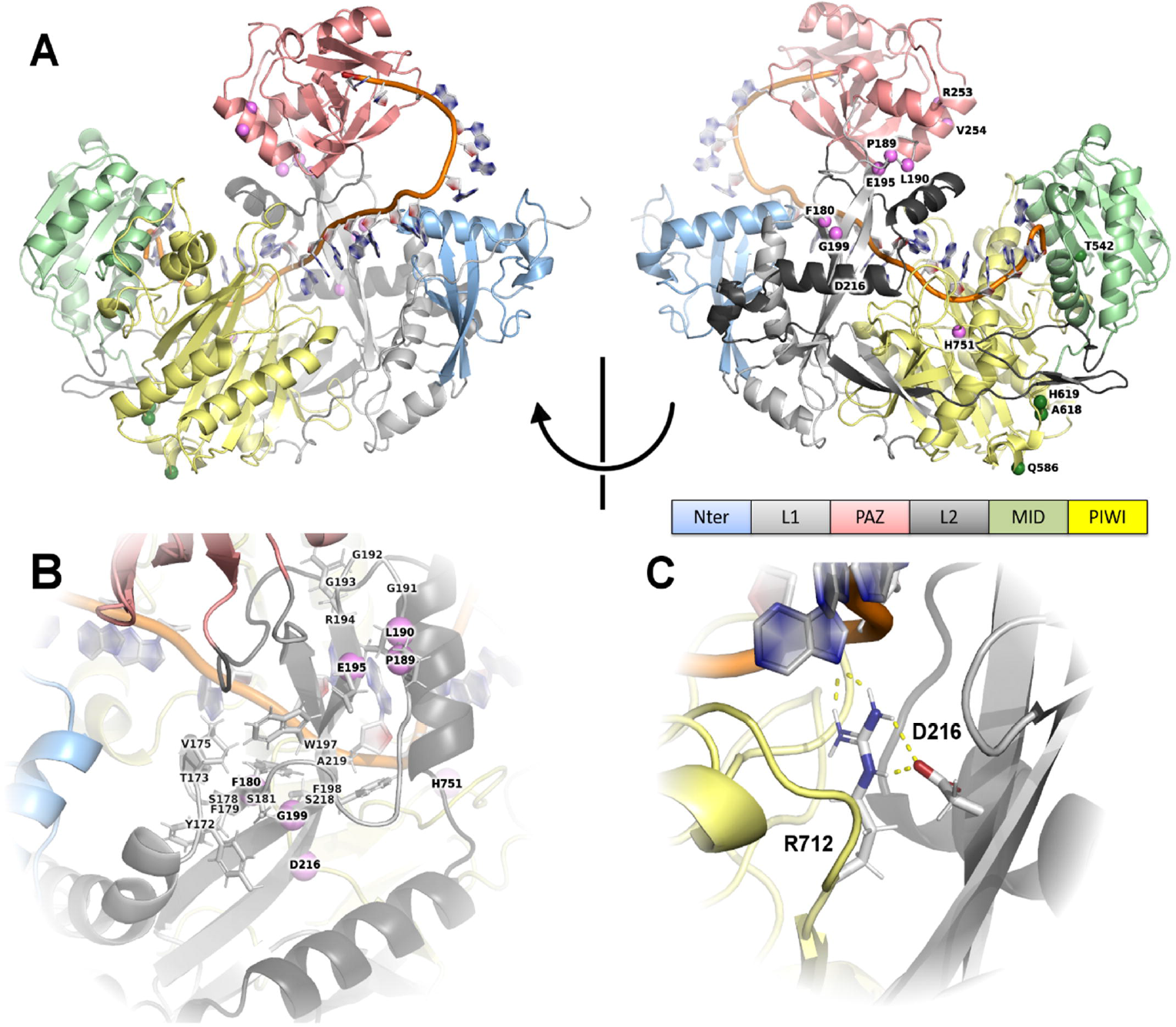
Facial characteristics of individuals with AGO1 variants. **(A)** Front faces, **(B)** profile faces (**C**) Face2Gene Facial analysis using Face2Gene Research application (FDNA Inc. Boston, MA) of unrelated individuals with *AGO1*-associated disorder (n = 14) compared to controls matched (n= 14) for sex, age, and ethnicity.

### Variants spatially cluster in AGO1

The AGO1 protein (NP_036331.1) contains five identified functional domains: two linker domains L1 (AA 173-225) and L2 (AA 372-418), one MID domain (AA 427-508), one PAZ domain (AA 226-368) and one PIWI domain (AA 515-816). The PAZ domain contains the RNA binding module that recognizes the 3′ end of both siRNA and miRNA. The PIWI domain contains the cleavage site that is inactive in AGO1 and is involved in protein/protein interactions, notably with DICER and GW182. The nonsynonymous variants cluster in 17 amino acid positions of the protein: eight are in the L1 binding domain (p.Phe180, p.Pro189, p.Leu190, p.Glu195, p.Gly199, p.Asp216, p.Val217_Ser218), four in the PAZ domain (p.Arg253, p.Val254, p.Pro324, p.Thr355, p.Gln358), two variants in the L2 domain (p.Glu376, p.Tyr418) and three in the PIWI domain (p.His751, p.Thr781, p.Ile797) **(Table 2)**.

The majority of 10 first collected variants affect amino acids (Phe180, Pro189, Leu190, Glu195, Gly199, Asp216, Arg253, Val254 and His751) clustered in 3D along the sides of L1 and PAZ domains facing one another in 3D, except for His751 **(Figure 3A)**. Indeed, within the PAZ domain, Arg253 is within an alpha-helix and makes specific contact with the last residue in the helix, Asp250, and the backbone of Arg284. Thus, polar contacts involving Arg253 are likely important for the organization and stability of the PAZ domain. The variant p.(Arg253His) may modify these interactions. The nearby variant p.(Val254Ile) is conservative, but valine has the lowest helical propensity of the small hydrophobic amino acids^35^. Thus, this variant could over-stabilize the helix and limit motion of the PAZ domain. Three critical amino acid positions are within the L1 domain and in proximity to the PAZ domain: Pro189, Leu190, and Glu195 **(Figure 3B)**. Pro189 is at the base of a long loop and the backbone geometry of proline may be necessary for limiting the loop’s mobility; p.(Pro189Leu) may thus alter loop dynamics. Leu190 packs within a hydrophobic interface made up of amino acids from L1 and L2; p.(Leu190Pro) or p.(Leu190Arg) may distort the interface between the linker domains. Glu195 is close in space to Trp197 and Lys224; p.(Glu195Lys) may repel these nearby amino acids and destabilize the L1 domain. Farther down the L1 domain, p.(Gly199Ser) is a recurrent variant (observed in four unrelated individuals and published once in another individual): the side chain of the introduced serine could clash with the side chain of Ser218 and introduce an unfavorable polar contact, likely destabilizing the L1 domain.

**Figure 3:**
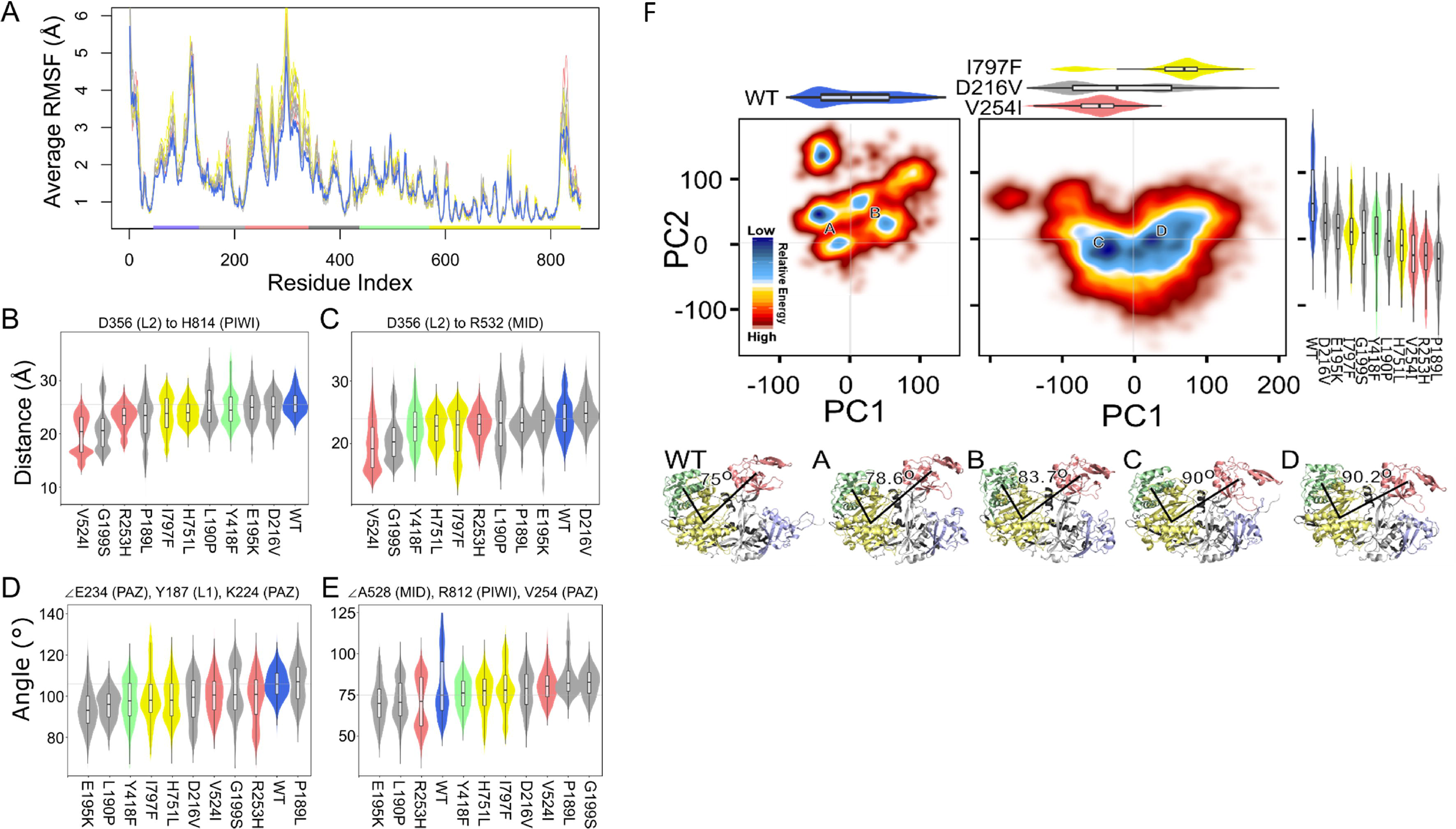
AGO1 structure and annotation of novel variants. **(A)** Protein structure of AGO1 colored by protein domain and with the sites of variants indicated by spheres. The linker domains, designated L1 and L2, are separated in sequence by the PAZ domain, but intertwine in 3D, forming common interfaces between the N-terminal, PAZ, and PIWI domains (**B)** Many of the variants of interest are within the first linker domain. This domain forms a narrow beta sheet with three strands. Variants occur in the middle of these strands at the closest point between domain L1 and the guide RNA backbone, and towards the end of L1 and near the PAZ domain interface (**C)** D216 is within L1 and makes specific contacts with R712 in the PIWI domain. R712 also interacts directly with the guide RNA.

Phe180 makes contacts across all three strands of the L1 beta sheet and its deletion would significantly alter the hydrophobic packing of the region. Asp216 makes a salt bridge with Arg712 in the PIWI domain **(Figure 3C)**. Arg712 also makes specific interactions with the guide RNA. Thus, the variant p.(Asp216Val) is likely to destabilize the interface between L1 and PIWI, and thereby indirectly alter interactions with the guide RNA. Finally, p.(His751Leu) is the only observed variant within the PIWI domain. His751 makes hydrogen bonds with the guide RNA and p.(His751Leu) is therefore likely to alter the stability of the interaction between AGO1 and the guide RNA.

### Variants are predicted to affect AGO1 dynamic conformational change

We used our molecular dynamic (MD) simulations to assess global and local changes to AGO1 in association with each genomic variant. We first assessed global changes to AGO1 conformation across our MD simulations using RMSD, RMSF and PC analysis (**Figure 4**). We found that genomic variants have the largest change in the patterns of mobility for the N-term and PAZ domains, and more modest mobility changes to the MID domain (**Figure 4A**). Because RMSF depends on how the trajectories are aligned, and we aligned based on the PIWI domain, we observed relatively small overall fluctuations of the PIWI domain. Therefore, patterns in RMSF indicated that the largest features in our simulations were domain-domain motions, and that these were modified by genomic variants.

Next, to assess domain-domain distance, we assessed linear distances from specific residues in the PIWI, MID, and linker regions. Certain PAZ and linker variants such as p.(Val254Ile), p.(Gly199Ser), p.(Arg253His), and p.(Pro189Leu) were associated with significantly shorter distances between the PIWI and linker **(Figure 4B)**. The same variants p.(Val254Ile) and p.(Gly199Ser) and additional ones p.(Tyr418Phe) and p.(His751Leu) were associated with significantly shorter distances between the MID and linker **(Figure 4C)**. Because the relative orientation of these domains is likely important for AGO1 function, we also assessed the angles between them. Genomic variants in multiple domains altered the orientation of PAZ with respect to linker **(Figure 4D)**, and the MID-PIWI-PAZ orientation **(Figure 4E)**. Thus, we identified additional and more specific changes in domain-domain orientations associated with genomic variants.

Because many conformational changes occur concurrently in our MD simulations, we next assessed all dynamics data together using PC analysis **(Figure 4F)**. We visualized the predominant dynamics of AGO1 using a free-energy landscape. We found that the WT protein sampled one side of the landscape, while variants had significant sampling of a different side. PC states are characterized by the relative orientations of the PAZ, Nterm, and MID domains (see **Supplemental Animation 1** for PC visualizations). Genomic variants had two effects on these predominant dynamics. First, they shift the average conformation to one that favors a wider angle of the PAZ-PIWI-MID domains. Second, the transition between conformations was blurred with a significant amount of time that the protein deviates from WT_like confirmations. This indicates that the genomic variants may change the nature of AGO1, leading to dysregulated dynamics. We explored the possibility for changes in internal distances as a summary of dysregulated dynamics (**Supplementary Figure S3**) and demonstrated that genomic variants were associated with lower coordination between domains. We anticipate that changes to domain coordination could be an additional and informative criterion for assessing multi-domain enzymes.

To further assess the intra-domain change associated with each amino acid variant, we calculated the per-domain RMSD **(Supplementary Figure S4)**. We found that amino acid changes throughout the structure could be associated with alterations of the same or distant domains. For example, p.(Leu190Pro) is within the linker domain and the conservative variant p.(Val254Ile) is within the PAZ domain but both lead to alterations in the internal organization of the linker and MID domains.

## DISCUSSION

This study presents the clinical and molecular characterization of a cohort of 28 individuals from 26 unrelated families with 15 unique heterozygous coding variants (13 missense and two in-frame deletion variants) in *AGO1*. This study establishes that *de novo* coding variants in *AGO1* are responsible for a form of NDD characterized by psychomotor delay, behavioral features and language impairment. Including the three previously reported variants, a total of 18 variants have been described in *AGO1* in a total of 33 individuals with NDD. Six variants are recurrent: p.(Gly199Ser), p.(Phe180del), p.(Leu190Pro), p.(Leu190Arg), p.(Val254Ile), p.(Glu376del), while the others are reported in only one family to date. All the variants lead to substitutions or deletions of one or two amino acids, but no truncating variant (nonsense, frameshift, or splicing variants leading to truncation) was observed in this cohort. Interestingly, four of the amino acids residues mutated in *AGO1*, Phe180, Leu190, Gly199 and Thr355, were also found mutated at the equivalent residues in *AGO2* (p.Phe182del, p.Leu192Pro, p.Gly201Cys or Val, p.Thr357Met) in ID patients **(Supplementary Figure S1)**. Others *AGO2* missense variants were reported in L1 domain (p.His203Gln), in helix-7 of the L2 domain (p.Met364Thr, p.Ala367Pro) involved in the proper positioning of the RNA guide, in the MID domain (p.Gly573Ser), and in PIWI domain (p.Gly733Arg, p.Cys751Tyr and p.Ser760Arg). The last three variants are located in a loop which binds in the minor groove of guide/target duplex^15^.

ID and language delay were reported in all affected while most of them displayed motor delay, seizures, autistic features, and behavior problems. No recurrent structural brain nor other organs malformation was noticeable and if individuals share some common facial traits, no typical specific gestalt was observed. Therefore, it appears that the NDD form caused by coding variants in *AGO1* is not a clearly recognizable entity. Moreover, the individuals present differences in the severity of the clinical manifestations, which do not appear to correlate with the nature or the location of the variant. The severity of ID, as well as growth parameters, was for instance highly variable among the three individuals with the p.(Phe180del) variant. We could therefore suggest that additional genetic or environmental factors might play a role in the phenotypic expressivity, modulating the effect of the *AGO1* variant. On the contrary, some specific traits seem to correlate with specific variants, such as hypothyroidism specifically reported in two individuals with the p.(Phe180del) variant, or status epilepticus in three individuals with p.(Gly199Ser) variant. A larger number of individuals would be necessary to confirm these observations. Individuals with *AGO1* or *AGO2* variants showed common clinical findings: ID with autistic features and aggressiveness, impaired speech development, motor delay, MRI anomalies and frequent gastrointestinal disorder or reflux. Minor additional clinical features are also reported in both cohorts: skeletal (clinodactyly or bradymetacarpy, scoliosis), vision problem (strabism, myopia/hyperopia, visual impairment), heart, dental or breathing anomalies, and also anxiety and sleeping disturbance. Frequency of such minor anomalies should be confirmed by futures studies. To note, we observed three monozygotic couples in *AGO1/AGO2* cohorts. This apparent excess of monozygotic twins (6/54 patients) have to be confirmed with future studies.

The 3D conformational model of AGO revealed a very flexible protein with two mobile domains, the L1 and L2 linker domains. The global structure contains four globular domains (PAZ, MID, PIWI and N terminal), and a deep cleft bordered by L1 and L2 linkers. L1 and PAZ domains seemed to be more flexible than the L2, PIWI and MID domain, as shown in **Figure 3**. The flexibility of the protein seems to be important to permit transitions between the different phases of the RISC process: sncRNA loading and processing, mRNA target clip, helper protein recruitment, and finally RISC complex release. Opening and closure of the PAZ/L1 jaw seems necessary for the proper function of AGO protein. Among the 18 *AGO1* variants reported to date the majority clustered within the L1, PAZ and PIWI domains. We have analyzed the potential effects of variants located in the L1 and L2 linker domains, as well as some of the variants located in the PAZ and PIWI domains, using structural biology as an interpretive framework. We considered the role of native residues within AGO1 and how they would be altered by the variants. Our assessment of the *AGO1* variants identified in individuals with NDD indicated that they were deeply buried in the cleft region and close to the RNA guide molecule, but did not affect the residues involved in RNA guide anchorage; in addition, dynamical simulations showed that these variants narrowed the angulation between the PAZ/L1 domains. These observations suggested these variants may hamper AGO1 flexibility during the different phases of the mRNA processing. Recently, the AGO1x protein isoform was described, product of the translational stop read-through of *AGO*1 transcripts induced by the expression of the Let7a miRNA. Ectopic expression of AGO1x acts as a negative competitor of the miRNA pathway^36^. A hypothesis would be that mutated AGO1 proteins may act as the AGO1x isoform through competitive inhibition of wild type AGO proteins. Functional studies will be necessary to investigate this hypothesis.

The absence of truncating *AGO1* variants in our cohort is surprising and is not in favor of a loss-of-function (LoF) as the unique molecular mechanism involved in *AGO1*-related ID. However, the observation of five children with large heterozygous deletions including *AGO1* together with *AGO3 (*and sometimes *AGO4* among other genes), and a deletion encompassing the *AGO2* fist 3 exons challenges this observation^13–15^. *AGO2* variant functional analysis revealed a complex cellular deregulation: a decrease in mRNA silencing was observed as expected for a LoF mechanism but with a variable impact, depending on the mRNA target and the tested *AGO2* variant. Reduced target release and reduced phosphorylation of the serine cluster at residues 824 to 834 in AGO2 was observed for most of the tested variants due to probable protein dynamics perturbation, likely leading to AGO2 deregulation and reduced functions ^15^.

A better understanding of how AGO1 functions in the brain will be essential to understanding how pathogenic variants in this gene cause cognitive impairment, developmental delays and behavioral manifestations including autistic traits. In the developing mouse brain, although it is ubiquitously expressed, *Ago1* was shown to be upregulated in neurogenic progenitors and mature neurons^37^. This expression pattern correlates with initial findings showing a role for human AGO1 in promoting differentiation after neuronal induction in a cellular model of neuroblastoma SH-SY-5Y cells^37^. Mouse models carrying homozygous inactivation of *Ago1* showed postnatal lethality as reported in IMPC (International Mouse Phenotyping Consortium, mousephenotype.org): only 9 % of homozygotes Ago1^tm1a/tm1a^ mice are obtained from heterozygous mating suggesting a prenatal lethality. The surviving homozygotes (both males and females) showed decreased body weight and increased anxiety (abnormal behavior in the open field). Pathogenic variants in several other genes encoding protein partners of AGO1/AGO2 involved in the regulation of mRNA decay and translation, such as CNOT1^38^, CNOT2^39^, CNOT3^40^ or DDX6^41^, have been reported to cause NDDs^42^. Apart from their role in post-transcriptional regulation, human AGO1 and AGO2 proteins are also involved in the regulation of transcription and splicing^9^. Therefore, alterations of these nuclear functions could also be involved in the pathophysiology of the neurodevelopmental disorders caused by coding variants in these genes.

In conclusion, this collaborative study, reporting molecular and clinical data from 26 families with heterozygous *de novo* coding variants in *AGO1*, confirms the involvement of rare coding variants in this gene in NDD. Future studies investigating transcriptional and posttranscriptional regulation defects or other dysfunction stemming from *AGO1* pathogenic variants will be important for elucidating the precise mechanisms of disease that may inform potential therapeutic strategies.

## ACKNOWLEDGEMENTS

We thank all the families for their participation in the study. We also thank the CREGEMES for its financial support. The authors want also to thank all the people from the Strasbourg Hospital molecular diagnostic lab, especially the team working on diagnosis of Intellectual Disability (Claire Feger, Elsa Nourisson, Céline Cuny, Sylvie Friedmann and Carmen Fruchart) and all the bioinformatics team. The authors want also to thank all the people from the Cambridge Broad Institute of MIT and Harvard, especially the team working on clinic and diagnostic of patients (Victoria de Menil, Elise Robinson, Patricia Kipkemoi and Anne O’Donnell Luria). We also thank the Centre National de Génotypage (Jean-François Deleuze, Robert Olaso, Anne Boland, and the technicians and bioinformaticians) for their participation in library preparation and DNA sequencing. We also thank the Center for Individualized Medicine at Mayo Clinic for their support. FL, GV and AG received funding from European Union and Région Normandie in the context of Recherche Innovation Normandie (RIN 2018). Europe gets involved in Normandie with the European Regional Development Fund (ERDF).

## WEB RESOURCES

CADD, https://cadd.gs.washington.edu/snv

GeneMatcher, https://genematcher.org/

GnomAD browser, https://gnomad.broadinstitute.org/

OMIM, https://www.omim.org/

SySID, https://sysid.cmbi.umcn.nl/

## FIGURE LEGENDS

**Figure 4: Simulations reveal changes in domain orientation associated with de novo missense AGO1 variants**

We used MD simulations to assess how the native structure of AGO1 would respond to the introduction of a subset of identified genomic variants. (**A)** Variability of each amino acid was quantified using RMSF after aligning each trajectory to the initial WT conformation of the PIWI domain and averaging across replicates. Domains are colored as in Figure 1 and each variant colored according to the domain it is within. (**B-E)** We monitored selected distances and angles as a simple way to assess conformational changes between the (**B)** linker and PIWI domains, (**C)** linker and MID domains, (**D)** the orientation of the PAZ domain, and (**E)** the openness of the RNA binding region. (**F**) We show the free energy landscape across molecular dynamics (MD) simulations as a color gradient from high-energy to low energy. Above, we show one-dimensional PC samplings as a combined violin and boxplot. The left-and right-hand panels summarize all data from the WT and from our novel variants, respectively. Selected variant’s PC1 sampling is shown above the panel, and all variant’s (that underwent MD) PC2 sampling to the side of the panel. Four regions of low energy are indicated by letters A-D. Representative images of AGO1 structure from the simulations taken from these points of low energy are shown below and labeled by the corresponding letters with the WT shown for comparison. To summarize one aspect of the difference between these four regions, we show the angle between the centers of mass of the MID, PIWI, and PAZ domains.

## SUPPLEMENTAL DATA

