## Supplementary material for "*De novo* coding variants in the *AGO1* gene cause a neurodevelopmental disorder with intellectual disability": Suplementaries

**Supplementary data**

**- Supplementary text**

**- Figure S1: Consequences of c.650-2A>G variant on splicing**

(**A**) Alamut view (Alamut® Visual, SOPHiA GENETICS, Lausanne, Switzerland) of the the c.650-2A>G variant, predicted to abolish the use of exon 6 acceptor splice site according to all the different splice programs tested (**B**) Blood mRNA was obtained from family F15 proband, reverse transcribed into cDNA and amplified by PCR. Sanger sequencing of PCR products shows that the *de novo* c.650-2A>G variant results in the use of a cryptic splice acceptor located upstream the canonical one which leads to a 6 nucleotide deletion (r.650_655del) in the mRNA transcripts (NM_012199.4) and a 2 amino acids deletion p.(Val217_Ser218) in the protein. Deleted nucleotides are boxed.

**- Figure S2: Sequence alignments for AGO1 ortholog proteins from eight species**

Sequence alignment between AGO1 and the different human AGO paralog proteins or the different AGO1 ortholog proteins from eight species. The amino acids in red correspond to the amino acids affected by variants identified in individuals with ID in AGO1 and in green those corresponding to variants found in the general population from gnomAD. Amino acids positions mutated in AGO1 but also affected by another amino acid change in gnomAD are highlighted in green. Amino acid residues mutated in individuals with ID in AGO2 are highlighted in yellow. Alignments were performed using CLUSTALW 2.1. AGO1: homo sapiens (Human) AGO1 (Q9UL18, 857aa; AGO3: homo sapiens (Human) AGO3 (Q9H9G7, 860aa); AGO4: homo sapiens (Human) AGO4(Q9HCK5, 861aa); AGO2: homo sapiens (Human) AGO2(Q9UKV8, 859aa); AGO1_PANTRO: Pan troglodytes (Chimpanzee) (A0A2I3SWL4,860aa); AGO1_MUSMUS: Mus musculus (Mouse) (Q8CJG1, 857aa); AGO1_XENLA: Xenopus laevis (African clawed frog) (A0A1L8H7N2, 857aa); AGO1_DROME: Drosophila melanogaster (Fruit fly) (Q32KD4, 984aa); AGO1_DANRER: Danio rerio (Zebrafish) (Brachydanio rerio) (K4I6K9, 858aa); ALG-1_CAEEL: Caenorhabditis elegans (Q3LTR7, 1010aa); AGO1_ARATH: Arabidopsis thaliana (Mouse-ear cress) (O04379, 1048aa); AGO1_SACPOM: Schizosaccharomyces pombe (strain 972 / ATCC 24843) (Fission yeast) (O74957,834aa).

**- Figure S3. Changes to internal distances can quantify average conformational change associated with genomic variants.** Enzymes such as AGO1 require coordination between domain for function. (**A**) We quantified coordination among AGO1 domains using the average change to internal distances. There are visually evident "blocks" in the data that indicate how motions of the enzyme are dominated by domain-domain motion. Variation within domains indicates movement within the respective domain. (**B**) We quantified the extent of coordination by the probability of residue pairs having a high or average change to internal distances. We compared the variants P189L and I797F, whose internal distance difference matrices are visualized in panels (**C**) and (**D**), respectively. We anticipate that changes to domain coordination could be an additional and informative criterion for assessing multi-domain enzymes.

**- Figure S4: Variants throughout AGO1 cause changes within multiple domains.**

Because PC motions were dominated by domain-domain motions, we used our molecular dynamic simulations for each variant to assess changes within each domain individually **(A)** We scored the statistical significance of each change from the WT using a resampling procedure (see Methods). We show a per-domain heatmap for if the within-domain conformational changes significantly differ from what is observed in the WT. Importantly, variants within a different domain may have effects on other domains **(B-F)** We show the distributions of each domain across variants, colored by the domain in which each variant resides.

**- Table S1: Clinical manifestations observed in individuals with variants in *AGO1***

SD, standard derivation; ND, not determined; twin dz, twin dizygotic; twin mz, twin monozygotic; perception deafness, percep. Deafness; a, status epilepticus; b, regression language; autistic behaviour: c, stereotypies with echolalia; yrs, years, MRI: Magnetic Resonance Imaging; CCA: corpus callosum agenesis; CCH: corpus callosum hypoplasia

**- Table S2. Comparison between clinical manifestations presented by individuals with missense variants in *AGO1* and those with deletions encompassing *AGO1-AGO3/AGO4* (from Tokita *et al.*, 2014)**

**- Video S1. Animation of the motions represented by PC1, 2, and 3 from the molecular dynamic simulations.** Domains are colored as in Figure 1. Blue spikes represent negative PC values, and red spikes positive PC values. The animation shows the PC motions concurrently and from three different synced points of view. Each PC is scaled to the same maximum distortion so that their relative shapes (extent of domain-domain motion) can be visually compared.

#### SUPPLEMENTARY TEXT

##### <u>Family 1</u>

###### Clinical description

The proband is a male born at full term via uncomplicated C-section without neonatal complications. Global developmental delay was noted in early childhood and he received an autism spectrum disorder diagnosis at age 4 years. He spoke his first word between 12-24 months; by age 10 years 9 months he spoke in short phrases. Spontaneous functional communication was limited, and echolalia was noted. Behavioral concerns have included intermittent aggression and impulsivity, though these have improved with age, and he has been diagnosed with ADHD and anxiety. EEG at age 7 was consistent with centrotemporal spike discharges during sleep; benign rolandic epilepsy was diagnosed and treated with Trileptal, which was successfully weaned at age 8 years. He has been followed by endocrinology for growth hormone deficiency and hypothyroidism. There is also a history of constipation. On physical exam, he was non-dysmorphic and muscle tone and strength were normal. He had a capillary hemangioma on his right knee. Full scale IQ, based on school evaluations at age 5 years, was 76 and he received learning support. His family history was negative for neurodevelopmental disorders, except for a maternal cousin reportedly diagnosed with Asperger syndrome. Chromosomal microarray was negative.

###### Sequencing method

Using genomic DNA from the proband and parents, the exonic regions and flanking splice junctions of the genome were captured using the IDT xGen Exome Research Panel v1.0. Massively parallel (NextGen) sequencing was done on an Illumina system with 100bp or greater paired-end reads. Reads were aligned to human genome build GRCh37/UCSC hg19, and analyzed for sequence variants using a custom-developed analysis tool. Additional sequencing technology and variant interpretation protocol has been previously described[1]. The general assertion criteria for variant classification are publicly available on the GeneDx ClinVar submission page (http://www.ncbi.nlm.nih.gov/clinvar/submitters/26957/). Exome sequencing (trio analysis) revealed a heterozygous, *de novo* in-frame deletion NM_012199.4:c.539_541delTCT, p.(Phe180del) in the *AGO1* gene. Additionally, a heterozygous, paternally inherited frameshift variant (NM_002693.2:c.1646delT) in *POLG* was reported. *POLG* is associated with recessive and dominant forms of mitochondrial disease which can manifest as progressive external ophthalmoplegia, seizure disorders, a range of neurological and muscular symptoms, and other symptoms. However, dominant forms of *POLG*-related disorders tend to be adult-onset. Given his father's negative clinical history and lack of a second variant in *trans*, this was considered unlikely to be causative

**Family 2**

**Clinical description**

The proband is a 3-years-old boy presenting with global developmental delay. There is no familial history except that the paternal grandfather had a stroke in his 30's. He was born at term and had a neonatal jaundice. He sat at 14 months and crawled at 19 months, stood at 21 months, and cruised around 2 years old. He had no speech at 2 years and communicated screaming. He had abnormal social interaction and intermittent eye contact. The proband presented with a truncal hypotonia more than appendicular. Plagiocephaly and brachycephaly were also observed at 2 months old. A notion of emesis after eating until 1 year old has been reported and constipation that had improved with age. He has eczema and café-au-lait spots, one on the head, one on the left wrist.

Standard karyotype and CMA (chromosomal microarray) were normal. Brain MRI (Magnetic Resonance Imaging) only showed increased extra-axial fluid.

**Sequencing method**

Genomic deoxyribonucleic acid (gDNA) is isolated from the patient and parent's whole blood. Full exome sequencing, bioinformatics analysis, filtering and manual review of alterations was performed as previously described Farwell, 2015[2] and Smith 2017[3] with Sanger sequencing of alterations with likely clinical relevance. Exome sequencing (trio analysis) revealed a heterozygous, *de novo* NM_012199.4: c.539_541delTCT p.(Phe180del) variant in *AGO1*.

**Family 3**

**Clinical description**

The proband is a 9 years old boy, the only child of non-consanguineous parents. There is no familial history, and the pregnancy was uneventful. I was born at 38,5 weeks with a weight at 3210g, a height of 48cm, and a head circumference of 35cm. He presented a severe global developmental delay, associated with autistic disorder. He could walk at 20 months old. He had not acquired verbal communication, but he can use PECS (Picture Exchange Communication System). He presented febrile seizures since the age of 5 years, treated by valproic acid. A gastrostomy was performed due to severe feeding difficulties, with gastro-eosophageal reflux and an eosinophilic esophagitis diagnosed at the age of 3 years. He also had unexplained periodic fever associated with cutaneous rash every 2 months, and a hypothyroidism under treatment. A left cryptorchidism was also surgically treated. Brain MRI was normal at the age of 2 years. Abdominal and cardiac ultrasounds were normal. Standard karyotype, array-CGH, and direct sequencing of *ARX* were normal.

**Sequencing method**

Library preparation, exome capture, sequencing and data analysis were performed by IntegraGen SA (Evry, France), using Twist Human Core Exome in-solution enrichment methodology (Twist Bioscience, San Fransisco, California), followed by paired-end 75 bases massively parallel sequencing on Illumina HiSeq4000 (Illumina, San Diego, California). Bases calling were performed with Illumina Real Time Analysis software (2.7.6), alignement on GRCh38 reference with Burrows-Wheeler Aligner, and variant call with GATK Haplotype Caller GVCF (3.7). Filtering and interpretation of variants were performed using Bench Lab NGS Cartagenia software (Agilent, version 5.0.2). Trio-based Exome sequencing found a *de novo AGO1* variant: NM_012199.4: c.539_541del, p.(Phe180del).

**Family 4**

**Clinical description**

This proband is an 8-year-old male born from non-consanguineous Caucasian parents. The pregnancy was characterized by intrauterine growth retardation. The eutocic delivery was carried out at 37.1 weeks. Hypoglycemia and respiratory problems were observed in the neonatal period, while growth delay was reported in both the neonatal period and during infancy. His infancy was characterized by global developmental delay; severe non-verbal autism and intellectual disability were diagnosed at 4 years. A bone age of 4 years was reported when the patient was 6 years old. At 8 years, the patient shows repetitive behaviors and stereotypic movements, ADHD, sleep disturbance, auto and hetero aggressive episodes. Hypoplasia of the corpus callosum and bilateral hippocampal dysplasia were detected by MRI. In addition, the patient presents myopia, lack of sphincter control and receives nutritional supplements given his low weight.

**Sequencing method**

The *AGO1* variant NM_012199.4: c.539_541del, p.(Phe180del) was detected through NGS on a MiSeq Illumina platform, using a custom panel that allows to sequence the coding sequence of 460 genes mostly related to dominant forms and X-linked forms. All the bioinformatic analysis was performed using an in-house pipeline. In summary: raw reads are mapped to the human reference genome (hg19) using BWA aligner and subsequently processed using the GATK pipeline in order to remove PCR duplicates and perform base quality score recalibration. Variants are identified using the Haplotype Caller algorithm from GATK. Identified variants are annotated using ANNOVAR and then filtered in order to remove: i) non-coding and non-splicing variants, ii) known likely benign/benign variants, iii) variants having a frequency >1/5000 in gnomAD, iv) and low quality variants. For the remaining variants several bioinformatics predictors are used to assess the deleteriousness of the variants. In particular in-frame variants are evaluated using PROVEAN. Inheritance pattern of the candidate variants is assessed through Sanger sequencing.

**Family 5**

**Clinical description**

The patient is a 6-year-old female with a history of autism spectrum disorder and developmental delays. The mother was gravida 2, para 2 and 29 years old at delivery. The father was age 26. The mother was on prenatal vitamins and had gestational diabetes controlled by diet during pregnancy. The first trimester screening and routine ultrasounds were normal. Her birth weight was 5 pounds and 14 ounces. She was discharged from the hospital after 3 days. At 24 months of age she was not walking independently and was receiving early intervention, physical therapy, occupational therapy, and speech therapy. She was seen by genetics at 24 months for developmental delays. At 24 months it was noted that she had muscular hypotonia, gross motor delays, hand flapping and repetitive behaviors. She was attempting to stand at 24 months of age but was not ambulating. At a follow up visit at 6 years of age she had made some improvements. She was walking and running, and had 75-100 words. A developmental pediatrics assessment at this time noted that she had made substantial progress but still met the clinic criteria for autism and required substantial support. Hypoplastic teeth and reduced enamel were noticed.

An acylcarnitine profile, total and free carnitine were normal. There was a normal microarray and the chromosomes were 46, XX normal female. Fragile X testing, and methylation testing for Prader-Willi and Angelman syndrome was negative.

**Sequencing method**

Standard laboratory methods were used for DNA preparation. For exome sequencing, DNA libraries were prepared using the Agilent SureSelectXT Clinical Research Exome kit (Illumina, San Diego, CA) and sequenced using NextSeq 550 (Illumina). Sequences were processed using DRAGEN Bio-IT Platform software (Illumina, San Diego, CA) and .vcf files were uploaded to emedgene (Tel Aviv-Yafo, ISreal) for filtering and subsequent analysis of variants of interest. The analysis revealed a heterozygous NM_012199.4: c.539_541del, p.(Phe180del) *de novo* variant in *AGO1*. Sanger sequencing of the *AGO1* gene was performed under standard PCR conditions to confirm the NGS finding.

**Family 6**

**Clinical description**

The male proband was 9 years old at his last evaluation with a clinical diagnosis of autism. He was born at term with a normal birth weight to a pregnancy complicated by preeclampsia. Heart decelerations occurred leading to vacuum assisted vaginal delivery. Initial Apgar scores were four to five at one minute and seven at five minutes. At ten minutes, he developed retractions. Therefore, he was treated with blow-by oxygen, and these breathing difficulties never recurred. Neurocognitive development was normal until 12-18 months of life where he experienced rapid loss of language skills. At the age of 13 years, he is non-verbal with severe aggressive behaviours and moderate intellectual disability. He has significant aggression requiring intense psychiatric support.

Previous negative genetic testing investigations include chromosomal microarray, MECP2 sequencing, Prader Willi/Angelman syndrome MLPA analysis, Fragile X molecular analysis, and extensive biochemical testing. Brain MRI with spectroscopy at 4 years of age was normal. The Autism Diagnostic Observation Schedule (ADOS) was measured at the age of 4, and the patient scored 15 consistent with a diagnosis of autism.

**Sequencing method**

Clinical exome sequencing for this individual was performed by the Department of Laboratory Medicine and Pathology at Mayo Clinic in Rochester, Minnesota, USA. Genomic DNA was extracted from blood from the proband, biological mother, and biological father; 97% of the exome was covered at a read depth of 20X or greater. The exome was captured utilizing a custom reagent developed by Mayo Clinic and Agilent Technologies, targeting 19,456 genes and 187,715 exons using 637,923 probes to capture a 54.1Mbp total region. Sequencing was performed on an Illumina HiSeq 2500 Next Generation sequencing instrument, using HapMap Sample NA12878 as an internal control. Paired-end 101 base-pair reads were aligned to a modified human reference genome (GRCh37/hg19) using Novoalign (Novocraft Technologies, Malaysia). Sequencing quality was evaluated using FastQC (www.bioinformatics.babraham.ac.uk/projects/fastqc/). All germline variants were jointly called through GATK Haplotype Caller and GenotypeGVCF[4]. Each variant was annotated using the BioR Toolkit[5] and subsequently evaluated for clinical relevance. The clinical exome sequencing identified a *de novo* variant in *AGO1*, NM_012199.4: c.566C>T, p.(Pro189Leu). Targeted Sanger sequencing was used to confirm the presence of both variants in the proband and presence in the heterozygous state in each parent.

#### <u>Family 7</u>

**Clinical description**

This female proband is now 10 years of age. She was the product of a known dichorionic pregnancy that was uneventful. Delivery occurred at 35+ weeks and at 1980g she was slightly smaller than her co-twin. There were no concerns in the immediate newborn period, however parents began having concerns by a few months of age. She was formally referred for a genetics assessment at age 16 months. She has a history of global developmental delay. She was sitting unsupported at 16 months of age and by age 2 she was cruising on furniture and doing some babbling. Shortly after age 3 she began walking but has exhibited an unsteady gait. Even at the latest assessment she was essentially non-verbal, and was demonstrating increasing behavioral frustration as a result. She demonstrates limited eye contact and requires constant attention. She has a history of seizures, which tend to emerge with fever but have been well controlled on clobazam (Frisium). Her EEG (electroencephalography) demonstrates the non-specific finding of slow background for age. Otherwise she has been generally healthy. The family history is unremarkable. Her parents are non-consanguineous and of South Asian descent. She has a twin sister and a younger sister, both of whom are healthy with no developmental or cognitive deficits. On examination at 9 years of age head circumference was 50cm (- 1.5 SD), height 114.9 cm (-3SD) and weight 19.1 kg (-2 to -3SD). She exhibits multiple distinctive features including brachycephaly, a broad forehead, apparent telecanthus and downturned mouth corners. She exhibits a mildly ataxic gait. Prior to diagnostic clinical trio whole exome sequencing, investigations including normal array CGH testing and molecular studies for Angelman syndrome. Investigations for inborn errors of metabolism including urine organic acids, urine and plasma amino acids, plasma acylcarnitine profile, very long chain fatty acids and 7 dehydrocholesterol studies were normal. Brain MRI revealed mild cerebral atrophy.

**Sequencing method**

Trio clinical exome sequencing was performed by the laboratory Blueprint Genetics. Median coverage was 254, 251 and 221 in the proband, her mother and father respectively (99.31, 99.31 and 99.48 % coverage at greater than 20X). While the clinical analysis is primarily focused on established disease genes, all de novo coding variants are considered as potential candidates. The only variant of interest identified in the proband was a *de novo*, novel missense change NM_012199.4: c.569T>G, p.(Leu190Arg) in *AGO1*, predicted as probably damaging, deleterious and disease causing by Polyphen, SIFT and MUTTASTER respectively.

**<u>Family 8</u>**

**Clinical description**

The proband, a girl, is the 6th child of healthy, consanguineous Moroccan parents. One brother died at the age of 2 weeks; he was diagnosed with tetralogy of Fallot. The pregnancy was uneventful. She was born at term after a vaginal delivery with vacuum-extraction. A shoulder dystocia was noted. She had a good start, her birth weight was 5.3 kg, her height was 52 cm. Her development was delayed; she sat unsupported at the age of 9 months, she walked at the age of 2 years, after the start of physiotherapy. She spoke her first words at the age of 3 to 4 years. A diagnosis of autism spectrum disorder was made. At the age of 5 years, she developed epilepsy, that was classified as lesional/ partial epilepsy with sudden sideward movements of the eyes, and jerks of the left arm and leg, as well as absence seizures. A brain MRI demonstrated a corpus callosum agenesis and colpocephaly. At the age of 10 years, she shows episodes of behavioral problems, with aggressive behavior, often associated with hand biting, and hyperactivity. She eats a lot, it is difficult to restrain her in her eating. She attends special education. She sleeps well but is awake early. An ophthalmological evaluation demonstrates hypermetropia, astigmatism and an alternating esotropia. Her hearing is normal. Clinical examination at the age of 12 years shows a height of 128 cm (<p3), a weight of 36.9 kg (p25-p50) and a head circumference of 54.9 cm (p75). She has a hyperlordosis without scoliosis. Her neurological examination was normal, besides a slightly broad-based gait and strabism. She has a round face with a high nasal bridge, deep-set eyes with narrow palpebral fissures and full eyelids. The eyebrows are bushy. Furthermore, a cupid-bow of the upper lip, slightly posteriorly rotated ears, a broad neck and a supernumerary nipple are noted. The hands and feet are broad. Genetic testing using SNP-array and a targeted gene panel for developmental brain abnormalities were normal.

**Sequencing method**

Exome sequencing on the patient and parents was performed with DNA samples obtained from leukocytes. Exome capture was carried out with the de SeqCAp EZ Human Exome v3 kit (Nimblegen, Roche) target enrichment kits, and sequencing was performed on HiSeq 1500 platforms (Illumina) through the use of paired end reads. WES data processing, sequence alignment to GRCh37 were performed using an in house developed pipeline according to international standards. Variant filtering and prioritization was performed using VariantDB[6]. This exome sequencing using a trio-analysis approach demonstrated the *de novo* NM_012199.4: c.569T>G, p.(Leu190Arg) variant in *AGO1*.

**Family 9**

Female probands are identical twins (monochorial diamniotic pregnancy) born at term (37 6/7 weeks) by caesarean for abnormal cardiac rhythm in tween 2. They are 3rd and 4th children of non-consanguineous healthy parents. Birth measurements were normal, tween 2 had an Apgar score of 2 at 1 minute and 6 at 5 minutes, she presented hypotonia. Neonatal auditive auto-emission in the right ear was absent for tween 1, following evaluations concluded in mild deafness. Tween 2 had bilateral profound deafness and required hearing aids in her first months of age. Neuropediatric evaluation at 9 months of age revealed hypotonia, absent ocular contact, nystagmus, strabismus, and poor growth (length/height at -2.5 – 3 SD) for both tweens associated with dysmorphism motivating referral to clinical genetics. Dysmorphic features included hight prominent forehead, anteverted nares, broad nasal tip, bushy eyebrows, thin vermillion of the upper lip, coarse features, and gingival overgrowth. They evolved with severe global developmental delay: ocular contact appeared at 3 years old, sitting was acquired after 18 months with kyphosis. Tween 2 presented pharmaco-resistant (LEVETIRACETAM, LAMOTRIGINE) light-sensitive epilepsy with inaugural status epilepticus at 3.5 years old and developmental regression, spasticity, poor contact, very few movement, and axial hypotonia (absence of sitting at 8.5 years old). Tween 1 presented stereotypic movements and developed epilepsy at 5.5 years old treated by LEVETIRACETAM. She could stand with aid by age 7.5. At last examination (8 years old), tween 1 could walk aided, cannot crawl on all fours, can stand sitting with kyphosis, can eat with a spoon but is mainly feeded by gastrostomy. Both tweens show profound ID with absent speech, feeding difficulties needing gastrostomy feeding, gastro-oesophageal reflux and frequent urinary tract infections.

Various metabolic and genetic explorations showed normal results: amino acid chromatography in plasma, organic acid chromatography in urine, CDG, lysosomal and peroxisomal explorations, mitochondrial explorations, standard karyotype, CGH-array, 12p tetrasomy, *EHMT1* sequencing, ID gene panel sequencing. Brain MRI found unspecific fronto-temporal atrophy, no calcifications were found in brain tomodensitometry, ERG and visual potential were normal, bone radiography showed anomalies secondary to hypomobility.

**Sequencing method**

Exome sequencing was performed on blood DNA of the first twin. DNA was captured using SureSelect Human All Exon V5 (Agilent, Santa Clara, CA, USA) enrichment kit and sequenced on HiSeq 4000 (Illumina, 2x100bp paired-end). Data processing, sequence alignment to GRCh37/hg19, variant calling were performed using the in-house bioinformatic pipeline STARK. SNV and CNV filtering and prioritization were performed using Varank[7] and AnnotSV[8] respectively. Parental DNA were analysed in pools of six individuals. Exome sequencing revealed a *de novo* missense variant NM_012199.4:c.569T>C p.(Leu190Pro) in *AGO1*

**Family 10**

**Clinical description**

This female proband was born at term (39 3/7 weeks) after an uneventful pregnancy. At the age of 2 months she was hospitalized because of an RS-bronchiolitis. In addition, she showed reflux with feeding problems. She started walking at the age of 20 months. Around the age of 3,5 years she spoke her first understandable words. Her IQ was tested at 71 (verbal 85, performal 62). She shows uncoordinated running and tires quickly. There is no record of sleep disturbance or behavioral concerns. Previous genetic analysis: SNP array analysis, metabolic investigations, DNA analysis of the *ARID1B* gene and 1143 genes involved in intellectual disability showed no abnormalities.

**Sequencing method**

Exome sequencing (trio analysis) was performed with enriched DNA (Agilent Sureselect Clinical Research Exome (CRE) Capture) run on the Hiseq 4000 platform (150bp paired-end, Illumina). A missense NM_012199.4: c.595G>A p.(Gly199Ser) variant was identified in *AGO1*.

**Family 11**

**Clinical description**

The proband is a 10-year-old girl with global developmental delay with no episodes of regression, hypotonia, intractable epilepsy with focal but also possible generalized seizures. Seizures began at 9 months in the setting of febrile illness. Onset of non-febrile seizure at 7 years old with possible right-side predominance. Recently, seizure semiology consists of right arm flexion then jerking and shaking of the right side, twisting of the right side of the face with drooling. The average duration is 3-5 minutes with a frequency of every 3-4 months. It is also not unusual to have 2 seizures in one day.

Brain MRI at 8 years: Smearing of gray-white matter differentiation at the depth of the left superior frontal sulcus, suspicious for a focal cortical dysplasia. EEG: frequent high amplitude right sided independent spike/wave activity, with few independent left sided spikes. Chromosomal Microarray, infantile epilepsy panel, *DEPDC5*, and *SCN1A* testing were normal.

**Sequencing method**

Using genomic DNA from the proband and parents, the exonic regions and flanking splice junctions of the genome were captured using the Clinical Research Exome kit (Agilent Technologies, Santa Clara, CA). Massively parallel (NextGen) sequencing was done on an Illumina system with 100bp or greater paired-end reads. Reads were aligned to human genome build GRCh37/UCSC hg19, and analyzed for sequence variants using a custom-developed analysis tool. Additional sequencing technology and variant interpretation protocol has been previously described[1] The general assertion criteria for variant classification are publicly available on the GeneDx ClinVar submission page (http://www.ncbi.nlm.nih.gov/clinvar/submitters/26957/). Exome sequencing revealed a NM_012199.4:c.595G>A p.(Gly199Ser) variant in *AGO1*.

#### Family 12

**Clinical description**

This female proband was born to an uncomplicated pregnancy and delivery. She had global developmental delay. She walked at the age of 24 months and said her first words at the age of 3 years. At the age of 12 years, she could speak in full sentences in both French and Arabic. She cannot read and can do only simple additions. Neuropsychology testing revealed a moderate intellectual disability. The patient presented with a status epilepticus with partial onset at the age of 8 years. She was treated with clobazam and remains well-controlled and seizure free. Her latest EGG at the age of 14 years revealed the presence of multifocal as well as generalized epileptic activity. The EEG also showed disturbance of the background rhythm. On exam at the age of 12.5 years, she had a height of 149.5 cm (25th percentile), weight of 54.3kg (75-90th percentile) and head circumference of 53.4 cm (50th percentile). She was not dysmorphic. Her general and neurologic examinations were unremarkable. Chromosomal microarray, karyotyping, Fragile X testing and metabolic studies (plasma lactate and ammonia measurements, plasma amino acids and urine organic acids chromatographies, isoelectric focusing of serum transferrin) were normal. Brain MRI performed at the age of 10 years revealed increased T2 and FLAIR signal in the frontal periventricular regions.

**Sequencing method**

A *de novo* NM_012199.4:c.595G>A p.(Gly199Ser) variant was identified in *AGO1*: Case 702.278. Hamdan *et al.*, 2014[9]

**Family 13**

**Clinical description**

The proband was born at 42 weeks with symmetrically large size. She had relatively slow linear growth in her first two years of life while maintaining appropriate weight for length. Subsequent growth has been steady, but she has remained short-statured (particularly when compared to her family). She has global delay without regression. She began speaking a few single words before two years of age and began using phrases by 3 years of age. She has hypotonia and did not walk independently until 3 years of age and continued to have a wide-based, unsteady gait for several years. Her attention span is limited, but she is not hyperactive and has a pleasant demeanour. She began having partial onset seizures at 22 months of age. She had inadequate control of seizures on oxcarbazepine followed by levetiracetam (with multiple episodes of status epilepticus). She has had better control on clobazam with supplemental non-prescription cannabidiol (0-2 seizures/year). Her seizures have often been associated with illness (not necessarily febrile).

MRI performed at 22 months of age showed a few small foci of increased signal in the subcortical white matter of the left frontal and occipital lobes seen on the flair images only. MRI at 8 years of age showed more prominent bilateral periventricular and subcortical T2/flair hyperintensities. A *de novo* c.595G>A p.(Gly199Ser) variant was identified in *AGO1*.

**Sequencing method**

Agilent SureSelect Clinical Research Exome kit targeting the exonic regions of genes from the Epilepsy Exome panel from extracted genomic DNA. Targeted regions are sequenced using

Illumina NextSeq technology with 150bp paired-end reads. Sanger sequencing was used for confirmation.

#### Family 14

##### Clinical description

The proband is a boy and was born to non-consanguineous parents in gestational week 41+2 with a weight of 3650 g, a length of 54 cm and a head circumference of 36 cm. During pregnancy, a maternal urinary infection was treated by antibiotics. The mother has scoliosis and cleft vertebrae, the father had speech delay in infancy. Two maternal half-siblings are healthy. He could sit at age 1 year and walk at age 20 months. He did not speak at age 3 years but had better perception and started to communicate by signs. At age 4 years his IQ was 61 (SON-R 21/2-7), and an autism-spectrum disorder was diagnosed (ADOS-2 and ADI-R). Age at last investigation was 5 years and 7 months. Height was 118 cm (75th centile), weight was 24 kg (90th to 97th centile), and head circumference was 51 cm (25th to 50th centile). At that time he spoke slurred in single words and 5-6 word sentences. He had a short attention span, was hyperactive and easily overstrained by large groups or surroundings. Parents reported that he was operated because of cryptorchidism, produced a lot of cerumen, and that he has difficulties to fall asleep.

Metabolic screening, EEG, cranial MRI had been normal. Also karyotyping, chromosomal microarray analysis and testing for Fragile-X and Angelman syndromes had been normal.

##### Sequencing method

Trio exome sequencing was performed after enrichment with Agilent SureSelect v6 on an Illumina HiSeq 2500 platform. Average coverage was >100fold and coverage >20 fold in >90% of enriched regions. Obtained variants were filtered for *de novo*, autosomal recessive and x-linked variants, and a *de novo* NM_012199.4:c.595G>A p.(Gly199Ser) variant was identified in *AGO1*.

#### Family 15

##### Clinical description

This proband was born at term with a weight appropriate for gestational age and good postnatal adaptation. During pregnancy the mother was treated with anitiinfectiva as an infection with toxoplasma gondii was suspected. After birth infection with toxoplasma could not be confirmed but he had a newborn sepsis with group B streptococci. As a baby he had major problems with self-regulation and was crying a lot. Between the age of 3 months and 3 years he had frequent cyanotic breath-holding spells; epileptic origin and iron deficiency were excluded. After pyelonephritis at an age of 1 year a double kidney was detected. His development was slow from early infancy. He was able to crawl at 15 months and to walk at 22 months, he spoke his first words with 3 5/12 years. He never gained urinary and fecal control. Intelligence testing at the age of 5 8/12 years revealed an IQ of 55 (SON-R 2 ½-7). He attends a school for children with special needs and is now able to copy letters and to read short words. He is also able to count and identify small quantities, but cannot perform additions or subtractions. He is friendly and communicative even though his attention span is short and he is very distractible. He doesn't like activities, he hasn't chosen himself. His motivation for school work is low.

His diagnostic work-up comprised conventional karyotyping, array-CGH and fragile X syndrome, as well as testing for variants *SLC1A2* and *PMM2*. MRI at an age of 1 year and a cardiologic work-up were normal. Neurophysiological examinations (VEP/AEP) at an age of 1 and 2 years were normal, as were metabolic diagnostics. Several EEGs (latest 01/19) have also showed normal results.

He has a normally developed sister and no other family members with developmental problems.

**Sequencing method**

Exome sequencing of the affected individual and his parents was performed using the Agilent SureSelect All Exon V6 kit (Agilent) for exome enrichment and a NovaSeq6000 (Illumina) platform for sequencing. Reads were aligned to the human reference genome (UCSC hg19), with Burrows-Wheeler Aligner (BWA,V.0.7.8-r455). The average coverage was >100x in all samples and more than 97% of the targeted regions were covered more than 20x. High quality indel and single nucleotide variant calling and annotation were performed using GATK v3.1 as well as SAMtools v.0.1.7 using standard filtering criteria. Copy number variations (CNVs) were detected with ExomeDepth and Pindel. Candidate genes were prioritized by searching for homozyogus and compound-heterozygous variants with a minor allele frequency < 0.1% in the in-house database comprising >20,000 exome datasets as well as de novo variants with a minor allele frequency < 0.001%. A *de novo* splice variant in AGO1 c.650-2A>G was identified.

#### Family 16

##### Clinical description

Not reported

##### Sequencing method

Targeted sequencing of coding sequences of 451 genes known to be involved in NDD was performed on proband's blood DNA. DNA was captured using custom Agilent Sureselect enrichment kit and sequenced on HiSeq 2500 (Illumina, 2x100bp paired-end). Data processing, sequence alignment to GRCh37/hg19, variant calling were performed using the in-house bioinformatic pipeline STARK. SNV and CNV filtering and prioritization were performed using Varank[7] and AnnotSV[8] respectively. Parental DNA were analysed in pools of 14 individuals. A *de novo* NM_012199.4: c.758G>A p.(Arg253His) variant was identified in *AGO1*.

#### Family 17

##### Clinical description

This individual is a 22 years old male, first child born to unrelated healthy parents after an uneventful pregnancy with normal parameters (weight 3400g, length 48 cm, head circumference 36.5 cm, Apgar scores 10/10). His younger brother was healthy too. At the age of 5 months, Patient MM had spasms (short tonic movements of limbs with upward deviation of the eyes), but EEG did not suggest epilepsy and anti-epileptic treatment was not considered at that time. These spasms stopped at the age of 18 months with a trial of sodium valproate. It is unclear, however, whether these spasms were epileptic in nature or not. Seizures did not recur afterwards. EEG performed at the age of 4 and 10 years displayed bitemporal spikes. Patient walked independently at 1 year old. At 3 years, he is described as isolated with poor communication skills (he spoke a few words), hyperactivity/attention disorder, eye avoidance and repetitive behaviours suggesting autism spectrum disorder. He underwent ABA and speech therapies for several years and attended a school for special needs. His interaction and learning skills improved with time, so that at the age of 5, he was able to write his name and started to speak understandable sentences at 6. At the age of 15, he was able to read with limited understanding, he had acquired autonomy in day-today life. At 18, he worked in a sheltered workshop and neuropsychological assessment showed mild intellectual deficiency (WAIS-IV VCI 63, WMI 71, POI 62, PSI 50). Physical examination at the age of 15 was normal, height was 163 cm (-0.5 SD), weight 52 kg (-0.5 SD), head circumference 59 cm (+3 SD). Brain MRI was normal.

**Sequencing method**

Trio exome sequencing was performed on a NextSeq 500 Sequencing System (Illumina, San Diego, CA), with a 2x 150bp high output sequencing kit after a 12-plex enrichment with SeqCap EZ MedExome kit (Roche, Basel, Switzerland), according to manufacturer's specifications. Sequence quality was assessed with FastQC 0.11.5, then the reads were mapped using BWA-MEM (version 0.7.13), sorted and indexed in a bam file (samtools 1.4.1), duplicates were flagged (sambamba 0.6.6), coverage was calculated (picard-tools 2.10.10). Variant calling was done with GATK 3.7 Haplotype Caller. Variants were then annotated with SnpEff 4.3, dbNSFP 2.9.3, gnomAD, ClinVar, HGMD, Variome Great Middle East and an internal database. Coverage for these samples was 93% at a 20x depth threshold. Exome sequencing revealed a *de novo* NM_012199.4:c.760G>A p.(Val254Ile) missense variant in *AGO1*.

**Family 18**

**Clinical description**

Pregnancy and birth were uneventful with normal parameters. The early development for this female proband was normal. At age 4 years, severe behavioural problems were onset: aggression, automutilation, tantrums and impulsive behavior: there was no formal diagnosis of adhd but impulsiveness and problems focusing are present. Motor development was normal and cognitive evaluation showed a borderline-mild ID. At age 4 years and 8 months onset of focal seizures with impaired awareness, initially provoked by fever. No other neurological or malformations were retrieved. Nonspecific facial dysmorphic features were noticed with epicanthic folds and a flat nasal bridge. Brain MRI was normal. Karyotyping and SNP-array were normal.

##### Sequencing method

Exome sequencing revealed a pathogenic variant at the heterozygous state in the *BBS7* gene and in the *ADAR* gene, in addition to the *de novo* NM_012199.4:c.760G>A p.(Val254Ile) variant in *AGO1*. These genes are associated with autosomal recessive disorders.

#### Family 19

##### Clinical description

The proband is a 2 year 4-month-old coloured Khoisan male, with Autism Spectrum Disorder (ASD) level 3, and global developmental delay. He was born at 38 weeks' gestation by caesarean section. His birth weight was 2885g, and APGAR scores of 8 and 9 at 1 and 5 minutes respectively. Postnatally there were no complications however he was noted to have bilateral club feet, and had serial castings done. His early developmental milestones were delayed; sitting at 8 months and taking his first independent steps at 21 months. His speech and language were delayed, babbled from 8 months and did not progress as expected. When seen at age 2 years 4 months, he was able to walk without support, run and climb on furniture. In terms of his fine motor ability he had an established pincer grasp, was able to scribble and make a tower of two blocks, however no bridge or train and did not explore objects. In terms of speech and language, he is essentially non-verbal and cannot identify any body parts. He cannot use simple gestures in communication like waving good-bye. He responds to his name but is unable to follow simple one step instructions. He is able to finger feed himself, tries to eat everything, but spills most of the food with a spoon. He is still in diapers (unable to control bladder or bowel), assists in dressing himself but always requires help. Occasionally displays violent behaviour through aggressive and irritable outbursts. Physical exam showed macrocephaly (80th percentile head circumference for age) and short stature (-4 SD below the mean for age). Ophthalmological and hearing tests were normal. No imaging was performed on this patient as part of his work-up. Immunizations were up to date and he was not receiving any chronic medication. There is no known family history of developmental disorders and no known consanguinity. He meets the following DMS criteria for ASD: Criteria of social communication and interaction- (i) Deficits in verbal communication (ii) Parallel play with peers, prefers to play alone (iii) Has reduced shared interests with peers (iv) Very rigid in play, can look at an ant and follow it for a long time. Criteria for Restrictive pattern of behaviour, interest and activities: (i) Likes flapping and flicking his fingers, body rocking and hand-flapping (ii) Loves to grind teeth (iii) Hypersensitivity to loud noise and (iv) Picky feeder.

**Sequencing method**

Whole exome sequencing and data processing were performed by the Genomics Platform at the Broad Institute of MIT and Harvard with an Illumina exome capture (38 Mb target) and sequenced (150 bp paired reads) to cover >90% of targets at 20x and a mean target coverage of >100x. Exome sequencing data was processed through a pipeline based on Picard and mapping done using the BWA aligner to the human genome build 37 (hg19). Variants were called using Genome Analysis Toolkit (GATK) HaplotypeCaller package version 3.4. A *de novo* c.971C>T, p.(Pro324Leu) variant in *AGO1* was identified at the heterozygous state.

**<u>Family 20</u>**

**Clinical description**

The proband is a 4 years old boy, the second child of non-consanguineous parents. There is no familial history, and the pregnancy was gestational diabetes. He was born at 39 GW with a weight at 3570g, a height of 49cm, and a head circumference of 35,5cm. He presented a severe global developmental delay, associated with autistic disorder. He could walk at 30 months old. He had not acquired verbal communication at 3years old ans uses pointing. A phimosis was surgically treated. Audition was normal. Array-CGH found a 15q13.3 duplication inherited from his mother.

**Sequencing method**

Library preparation, exome capture, sequencing and data analysis were performed by IntegraGen SA (Evry, France), using Twist Human Core Exome in-solution enrichment methodology (Twist Bioscience, San Fransisco, California), followed by paired-end 75 bases massively parallel sequencing on Illumina HiSeq4000 (Illumina, San Diego, California). Bases calling were performed with Illumina Real Time Analysis software (2.7.6), alignement on GRCh38 reference with Burrows-Wheeler Aligner, and variant call with GATK Haplotype Caller GVCF (3.7). Filtering and interpretation of variants were performed using Bench Lab

NGS Cartagenia software (Agilent, version 5.0.2). The trio-based exome sequencing identified a *de novo* variant in *AGO1* NM_012199.4: c.1073A>G p.(Gln358Arg).

**Family 21**

**Clinical description**

The proband is a 3 year old white male of Welsh and Eastern European ancestry with developmental delay associated with postnatal microcephaly, hypotonia, dysmorphic features, and complex partial seizures. He was born after a full-term uncomplicated pregnancy by caesarean section with birth weight of 3.6 kg. There were no neonatal problems, but he was noted to be hypotonic from infancy. At birth, his head circumference was 33 cm (approximately 10th percentile for term infant), but his OFC gradually tracked below the 5th percentile for his age. His early developmental milestones were delayed; at age 11 months, he was not sitting up, crawling, holding a bottle, or saying any words. Brain MRI done at 10 months of age was unremarkable. When seen at age 3, he was able to walk but not run. He could jump and climbs stairs on his knees. He did not point or use gestures and he did not speak words but did make noises to indicate that he was hungry or tired. He appeared to understand simple directions. Exam showed bilateral epicanthal folds with narrow palpebral fissures, small ears with overfolded thick helices, flat elongated philtrum, thin vermillion border to upper lip, and microcephaly. He was hypotonic with normal deep tendon reflexes and was non-verbal. Ophthalmological and hearing tests were normal. He was treated with antiseizure medications for generalized seizures and underwent corpus callosotomy at age 7 due to intractable seizures. At age 8 he had a cardiac arrest in the setting of a respiratory infection; he was resuscitated and subsequently found by magnetic resonance imaging to have ischemic brain injury. There is no known family history of developmental problems. Father had been treated for learning disabilities as a child. He has a younger full sister who is well and older two half-brothers who are well. There is no known consanguinity.

Array CGH and serum lactate and pyruvate were normal, as well as genetic testing for *UBE3A, MECP2, TCF4, SLC9A6, FOXG1, CDKL5, FLNA, ASPM, CDK5RAP2, CENPJ, MCPH1, STIL,* and *CEP152*, and methylation analysis for Prader Willi and Angelman syndromes.

**Sequencing method**

DNA was isolated from patient and parent blood using QIAsymphony (Qiagen). Sequencing libraries were constructed from patient whole blood genomic DNA using HudsonAlpha Clinical Sequencing Lab's custom whole genome library preparation protocol. Patient DNA was sequenced on the Illumina HiSeq X sequencer. DNA library fragments were sequeneced from both ends (paired) with a read length of 150 base pairs. Patient genomes were sequenced at an approximate depth of 30X, with at least 80% of base positions reaching 20X coverage. Reads were aligned and variants called according to standard protocols[10,11]. A robust relationship inference algorithm (KING) was used to confirm familial relationships[12]. Sequenced variants were loaded into a custom software analysis application called Codicem for interpretation and variant pathogenicity was determined using ACMG criteria. Variants were Sanger confirmed by an external CAP/CLIA laboratory (EGL Genetics, Tucker, GA[MT1]). Exome sequencing revealed a paternally inherited variant in *ANKRD11* (c.4250A>G, p.Asp1417Gly), but given the paternal inheritance and lack of features of KBG syndrome, this was deemed to be non-diagnostic. Subsequent whole genome sequencing revealed three variants: 1) *SPG11:*c.5148dupA - judged to be pathogenic and not inherited from mother, but DNA was not tested from father; associated with hereditary spastic paraplegia but not felt to explain proband's microcephaly and neurodevelopmental disorder; 2) *SYNJ1:*c.3364+2T>G – likely pathogenic but maternally inherited; but not known to be associated with disease; 3) *AGO1*: 1 amino acid deletion, NM_012199.4:c.1126_1128delGAG, p.(Glu376del), noted to be *de novo.*

**Family 22**

##### Clinical description

Pregnancy was uneventful and birth was achieved at 41 weeks of gestation. Microcephaly (-3 SD) was noticed at birth with diffuse mild axial hypotonia with normal reflexes. Postnatal mild microcephaly, hirsute forehead, mild to moderate ID with speech delay were noticed. Mild spastic diplegia was also reported. Her coordination was abnormal: specific coordination testing such as rapid alternating movements was not possible, but by observation she did have excessive movements typical of a dyskinesia. Autistic features were observed with many stereotyped movements. Finally, bilateral moderate metatarsus adductus and lymphedema of the dorsa of both feet were also present.

##### Sequencing method

Not reported

**Family 23**

**Clinical description**

The proband is a 9-year-old boy born full term after a reported complicated prenatal history with illicit drug exposures. His birth weight was 3.51kg but further prenatal and perinatal details are not available as the child was adopted very early on in life. Family history is unknown. He walked at 18 months and with first words at 20 months of age. He received the diagnosis of autism at age 4 years. During the subsequent years he has required ongoing mental health care for his history of ADHD, oppositional defiant disorder, sensory processing disorder, and anxiety. He has required a variety of psychotropic medications for his behavioral issues and ongoing severe sleeping difficulties. His full scare IQ at age 9 of 78. On examination with no major dysmorphic features or neurological exam abnormalities A brain MRI scan and an ambulatory EEG performed at 7 years of age were unremarkable. Molecular cytogenetic studies (Affymetrix Cytoscan Dx) showed a benign CNV with an 81kb deletion at 6p25.3 (264,755-346,084).

**Sequencing method**

Using genomic DNA from the proband, the exonic regions and flanking splice junctions of the genome were captured using the IDT xGen Exome Research Panel v1.0. Massively parallel (NextGen) sequencing was done on an Illumina system with 100bp or greater paired-end reads. Reads were aligned to human genome build GRCh37/UCSC hg19, and analyzed for sequence variants using a custom-developed analysis tool. Additional sequencing technology and variant interpretation protocol has been previously described[1]. The general assertion criteria for variant classification are publicly available on the GeneDx ClinVar submission page (http://www.ncbi.nlm.nih.gov/clinvar/submitters/26957/). This analysis revealed a heterozygous in *AGO1* variant, NM_012199.4: c.1253A>T, p.Tyr418Phe).

**Family 24**

**Clinical description**

Twin probands A and B are two 21 years-old girls. They were born, at a gestational age of 33 weeks. Their parents are healthy nonconsanguineous parents. For the first twin, the birth weight was 1.980 kg, the birth length was 41 cm. For the second twin, the birth weight was 2.200 kg, the birth length was 43 cm. they both had a microcephaly, the head circumference was 50 cm (< -3 SD). They first presented an abnormal behavior with anxiety, aggressivity and oppositional-defiant behavior associated sleep troubles. They also had a motor and speech developmental delay. They walked alone at 24 months and they spoke at the age of 3.5. Intellectual disability was diagnosed in twin, in A as mild and in B as severe. On clinical examination, there was a microcephaly (-2.5 SD) and a nonspecific facial dysmorphic feature with a hypoplasia of the nose root, a columella forward, large ears normally hemmed, and an elongated face, associated with a camptodactyly (4th and 5th fingers) appeared around the age of 14.

Because of this clinic, a cerebral RMI was performed at the age of 11. It showed a very little atrophy of the upper vermis with hypoplasia of the lower vermis for A and a slight atrophy of the cerebellar and the vermis hemispheres without brain upstairs anomaly for B. Genetic tests were screening: a karyotype, CGHarray and Rett syndrome genes, MLPA, were normal.

**Sequencing method**

Targeted sequencing of coding sequences of 456 genes known to be involved in NDD was performed on blood DNA of the first twin. DNA was captured using Agilent Sureselect enrichment kit and sequenced on HiSeq 4000 (Illumina, 2x100bp paired-end). Data processing, sequence alignment to GRCh37/hg19, variant calling were performed using the in-house bioinformatic pipeline STARK. SNV and CNV filtering and prioritization were performed using Varank[7] and AnnotSV[8] respectively. Parental DNA were analysed in pools of 14 individuals. This targeted sequencing revealed a *de novo* missense variant in *AGO1* gene, NM_012199.4: c.2252A>T p.(His751Leu).

<u>**Family 25**</u>

**Clinical description**

The patient is a product of a 36 weeks 2 days twin gestation. Prenatally, he was exposed to alcohol, cocaine, and tobacco. Neonatal period was complicated by congenital HIV and HSV infections, thrombocytopenia, hypoglycemia, anemia, PFO, and GERD. Birth weight was

1.76kg. He required admission to the NICU for 5 weeks, during which time he underwent gastrostomy for GERD and failure to thrive. He had bilateral inguinal hernia repair at age 5 months. He had nystagmus and esotropia requiring bilateral rectus recession at ages 22 months and 5 years 5 months. He was prescribed glasses at age 7 years to assist with midline vision. At age 7 years, he retained is gastrostomy tube and required some gastrostomy tube feeds as he recently developed issues with swallowing. He had concerns for urinary incontinence, and heel cord tightness. At 7 years, his height was 111.90 cm (third percentile for age), weight was 20.80kg (twenty-second percentile for age), and OFC was 50cm (sixth percentile for age). He has upslanting palpebral fissures with epicanthal folds, prominent ears, a flat nasal bridge with small, upturned nasal tip, and full lips. The patient demonstrated motor delays, learning disorder, and behavioral issues. He sat independently at 11 months and walked at 18 months. He began saying single words at 9 months and combined two words at age 14 months. By 6 years 4 months, he was not able to recognize letters and had poor handwriting, but he was able to correctly identify colors, shapes, and his name when written. Though he was toilet trained by age 6 years 4 months, he did have frequent accidents. By age 7 years, he was not able to write his name and could recognize only some letters. Full scale IQ testing revealed IQ of 77 on the Stanford-Binet Intelligence Scales, Fifth Edition. Knowledge was identified as the highest Factor Index score in the patient's profile, while working memory was his poorest area of performance on educational testing. He has been diagnosed with learning disorder, unspecified behavior or impulse control disorder, anxiety, and ADHD. Family history information is limited as the patient is adopted. However, he was adopted with a twin sister who had speech delay and died at age 17 months from Hemophagocytic Lymphohistiocytosis.

Head MRI at age 5 months was normal. The patient had a normal chromosomal microarray via a custom designed ISCA 8x60k oligo-array at age 5 months. He then received genome sequencing given his persistent failure to thrive, short stature, developmental delays, learning disability, and behavioral issues.

**Sequencing information:**

DNA was isolated from patient and parent blood using QIAsymphony (Qiagen). Sequencing libraries were constructed from patient whole blood genomic DNA using HudsonAlpha Clinical Sequencing Lab’s custom whole genome library preparation protocol. Patient DNA was sequenced on the Illumina HiSeq X sequencer. DNA library fragments were sequeneced from both ends (paired) with a read length of 150 base pairs. Patient genomes were sequenced at an approximate depth of 30X, with at least 80% of base positions reaching 20X coverage. Reads were aligned and variants called according to standard protocols[10,11]. A robust relationship inference algorithm (KING) was used to confirm familial relationships[12]. Sequenced variants were loaded into a custom software analysis application called Codicem for interpretation and variant pathogenicity was determined using ACMG criteria. Variants were Sanger confirmed by an external CAP/CLIA laboratory (EGL Genetics, Tucker, GA[MT1] ). This genome sequencing (proband only) identified a heterozygous variant in *AGO1* NM_012199.4: c.2342C>T, p.(Thr781Met). Targeted testing for this AGO1 variant was performed at a CAP/CLIA certified lab and confirmed the presence of this variant.

**Family 26**

##### Clinical description

This 9-year-old boy was born after uneventful pregnancy with normal birth growth parameters. He initially showed hypotonia and a global psychomotor development. He walked at 18 months after intensive physiotherapy. His language was mildly delayed; he actually shows relative strength in language skills. He has major learning difficulties: his IQ measurements (WISC-V) show heterogeneous scores (VCI 84, VSI 61, FRI 55, WMI 69). He is treated for ADHD and manifests fine motricity impairment, as well as anxiety and behavioural trouble (strong need for routine; claustrophobia). There is no ASD or epilepsy. His brain MRI is normal. He has been operated for divergent strabismus and wears glasses for hypermetropia. He has primary nocturnal enuresis, and nocturnal terrors for which he receives Melatonin. Physically, the patient has no dysmorphic features, but exhibits relative macrocephaly and a slender build (poor growth compared to his siblings).

##### Sequencing method

**Figure S1.**

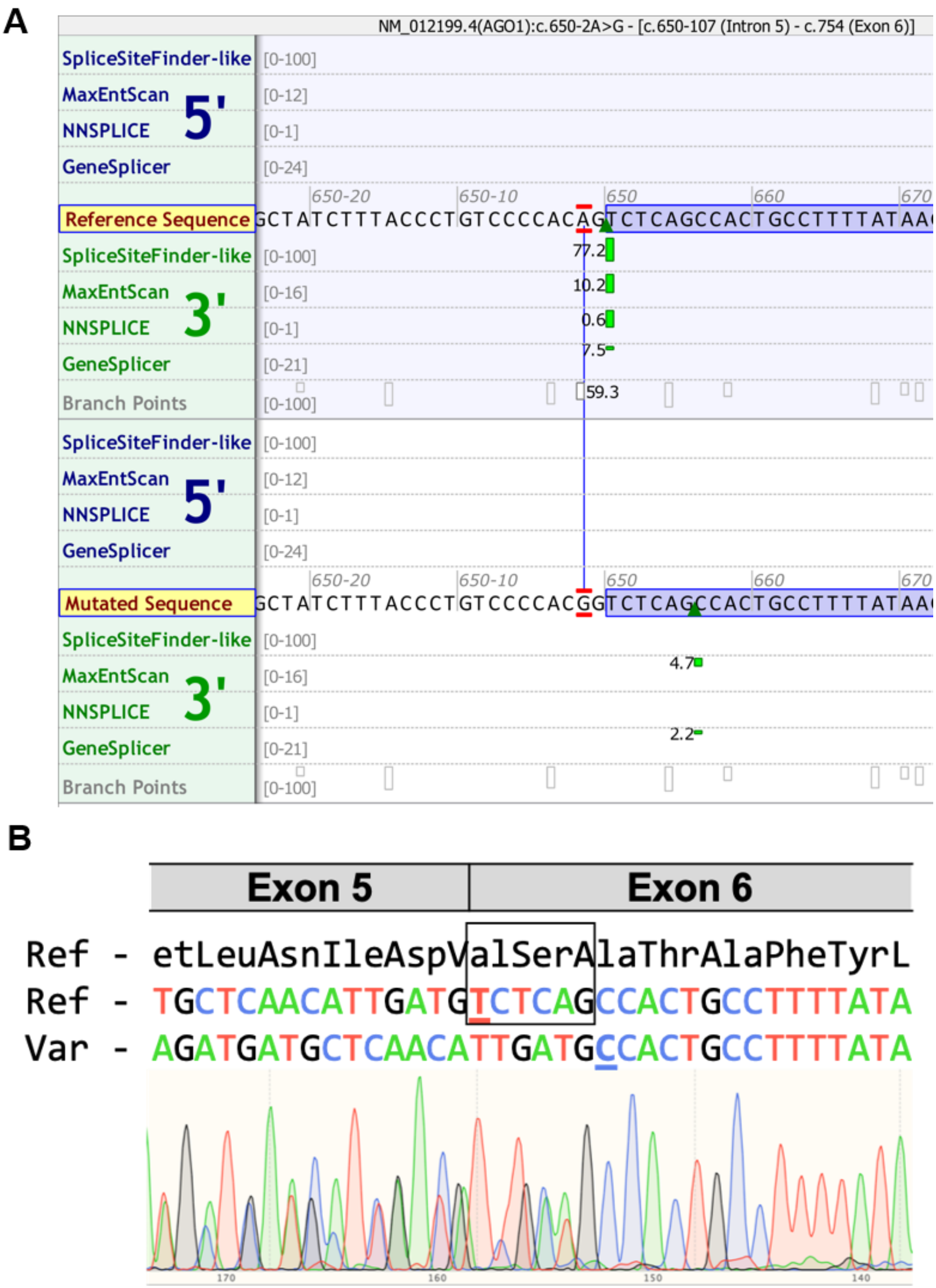
A
NM_012199.4(AGO1):c.650-2A>G - [c.650-107 (Intron 5) - c.754 (Exon 6)]
SpliceSiteFinder-like
MaxEntScan
NNSPLICE
GeneSplicer
5'
3'
Reference Sequence
Mutated Sequence
Branch Points
[0-100]
[0-12]
[0-1]
[0-24]
[0-16]
[0-21]
650-20
650-10
650
660
670
77.2
10.2
0.6
7.5
59.3
4.7
2.2
B
Exon 5
Exon 6
Ref - etLeuAsnIleAspValSerAlaThrAlaPheTyrL
Ref - TGCTCAACATTGATGTCTCAGCCACTGCCTTTTATA
Var - AGATGATGCTCAACATTGATGCCACTGCCTTTTATA
170
160
150
140

#### Figure S2

```
AGO1            ------------------------------------------------------------
AGO3            ------------------------------------------------------------
AGO4            ------------------------------------------------------------
AGO2            ------------------------------------------------------------

AGO1_PANTRO     ------------------------------------------------------------
AGO1_MUSMUS     ------------------------------------------------------------
AGO1_XENLA      ------------------------------------------------------------
AGO1_DROME      ------------------------------------------MSTERELAPGGPAQLHPH
AGO1_DANRER     ------------------------------------------------------------
ALG-1_CAEEL     -------------------------------SNVASNDPMSGGPQYLPGVMNSTIQQQPQ
AGO1_ARATH      MVRKRRTDAPSEGGEGSGSREAGPVSGGGRGSQRGGFQQGGGQHQGGRGYTPQPQQGGRG
AGO1_SACPOM     ------------------------------------------------------------


AGO1            ------------------------------------------------------------
AGO3            ------------------------------------------------------------
AGO4            ------------------------------------------------------------
AGO2            ------------------------------------------------------------

AGO1_PANTRO     ------------------------------------------------------------
AGO1_MUSMUS     ------------------------------------------------------------
AGO1_XENLA      ------------------------------------------------------------
AGO1_DROME      TLPLTFPDLQMTSTVGIIGKVYESQWTPSPTRPQSPSQAQTSFDTLTSPPAPGSSVNPTA
AGO1_DANRER     ------------------------------------------------------------
ALG-1_CAEEL     SATSSFLPSGPISSTSTSSQVVPTSGATQQPPFPSAQAAASTALQNDLEEIFNSPPTQPQ
AGO1_ARATH      GRGYGQPPQQQQQYGGPQEYQGRGRGGPPHQGGRGGYGGGRGGGPSSGPPQRQSVPELHQ
AGO1_SACPOM     ------------------------------------------------------------


AGO1            ------------------------------------MEAGPSGAAAGAYLPPLQQVFQAP 24
AGO3            ------------------------------------------MEIGSAGPAGAQPLLMVP
AGO4            --------------------------------------------MEALGPGPPASLFQPP
AGO2            ----------------------------------MYSGAGPALAPPAPPPPIQGYAFKPP

AGO1_PANTRO     ---------------------------------GDGGGGPLCLVAAGAYLPPLQQVFQAP
AGO1_MUSMUS     ------------------------------------MEAGPSGAAAGAYLPPLQQVFQAP
AGO1_XENLA      ------------------------------------MEAGPSSAAPSGLVPPLQQVFQAP
AGO1_DROME      VTSPSAQNVAAG--GATVAGAAATAAQVASALGATTGSVTPAIATATPATQPDMPVFTCP
AGO1_DANRER     -----------------------------------MEPGPSGAVPVGPYPPPLQQVFQAP
ALG-1_CAEEL     TFSDVPQRQAGS--LAPGVPIGNTSVSIGEPANTLGGGLPGGAPGQLPGGNQSGIQFQCP
AGO1_ARATH      ATSPTYQAVSSQPTLSEVSPTQVPEPTVLAQQFEQLSVEQGAPSQAIQPIPSSSKAFKFP
AGO1_SACPOM     --------------------------------------------------MSYKPSSEIA
                                                                  .

AGO1            RRPGIGTVGKPIKLLANYFEVD-IPKIDVYHYEVDIKPDKCPRRVNREVVEYMVQHFKPQ 83
AGO3            RRPGYGTMGKPIKLLANCFQVE-IPKIDVYLYEVDIKPDKCPRRVNREVVDSMVQHFKVT
AGO4            RRPGLGTVGKPIRLLANHFQVQ-IPKIDVYHYDVDIKPEKRPRRVNREVVDTMVRHFKMQ
AGO2            PRPDFGTSGRTIKLQANFFEMD-IPKIDIYHYELDIKPEKCPRRVNREIVEHMVQHFKTQ

AGO1_PANTRO     RRPGIGTVGKPIKLLANYFEVD-IPKIDVYHYEVDIKPDKCPRRVNREVVEYMVQHFKPQ
AGO1_MUSMUS     RRPGIGTVGKPIKLLANYFEVD-IPKIDVYHYEVDIKPDKCPRRVNREVVEYMVQHFKPQ
AGO1_XENLA      RRPGIGTVGKPIKLLANYFEVD-IPKIDVYHYEVDIKPDKCPRRVNREVVEYMVQHFKPQ
AGO1_DROME      RRPNLGREGRPIVLRANHFQVT-MPRGYVHHYDINIQPDKCPRKVNREIIETMVHAYS-K
AGO1_DANRER     RRPGMGTVGKPIKLLANYFEVE-IPKMDVFHYEVDIKPDKCPRRVNREVVEYMVQHFKPQ
ALG-1_CAEEL     RRPNHGVEGRSILLRANHFAVR-IPGGTIQHYQVDVTPDKCPRRVNREIISCLISAFSKY
AGO1_ARATH      MRPGKGQSGKRCIVKANHFFAE-LPDKDLHHYDVTITPEVTSRGVNRAVMKQLVDNYRDS
AGO1_SACPOM     LRPGYGGLGKQITLKANFFQIISLPNETINQYHVIVGDGSRVPRKQSQLIWNSKEVKQYF
                 **. *  *:   : ** *    :*   :  *.: :         :  ::
```

```
AGO1            IFGDRKPVYDGKKNIYTVTALPIGNERVDFEVTIPGEG-----KDRIFKVSIKWLAIVSW 138
AGO3            IFGDRRPVYDGKRSLYTANPLPVATTGVDLDVTLPGEGG----KDRPFKVSIKFVSRVSW
AGO4            IFGDRQPGYDGKRNMYTAHPLPIGRDRVDMEVTLPGEG-----KDQTFKVSVQWVSVVSL
AGO2            IFGDRKPVFDGRKNLYTAMPLPIGRDKVELEVTLPGEG-----KDRIFKVSIKWVSCVSL

AGO1_PANTRO     IFGDRKPVYDGKKNIYTVTALPIGNERVRLGVTIPGEG-----KDRIFKVSIKWLAIVSW
AGO1_MUSMUS     IFGDRKPVYDGKKNIYTVTALPIGNERVDFEVTIPGEG-----KDRIFKVSIKWLAIVSW
AGO1_XENLA      IFGDRKPVYDGKKNIYTVTTLPIGHERVDFEVTIPGEG-----KDRIFKVSIKWVAVVSW
AGO1_DROME      IFGVLKPVFDGRNNLYTRDPLPIGNERLELEVTLPGEG-----KDRIFRVTIKWQAQVSL
AGO1_DANRER     LFGDRKPVYDGKKNIYTVLALPIGSEKVDFEVTIPGEG-----KDRIFKVSIRWLAKVSW
ALG-1_CAEEL     FTN-IRPVYDGKRNMYTREPLPIGRERMDFDVTLPGDSA----VERQFSVSLKWVGQVSL
AGO1_ARATH      HLGSRLPAYDGRKSLYTAGPLPFNSKEFRINLLDEEVGAGGQRREREFKVVIKLVARADL
AGO1_SACPOM     GSSWMNSVYDGRSMCWSKGDIADGTIKVNIGSES---------HPREIEFSIQKSSKINL
                  .   . :**:   ::   :.     . :               : : . ::  .  .


AGO1            RMLHEALV--------SGQIPVPLESVQALDVAMRHLASMRYTPVGRSFFSPPEG----- 185
AGO3            HLLHEVLTGRTLPEPLELDKPISTNPVHAVDVVLRHLPSMKYTPVGRSFFSAPEG-----
AGO4            QLLLEALAGHLN--------EVPDDSVQALDVITRHLPSMRYTPVGRSFFSPPEG-----
AGO2            QALHDALSG--------RLPSVPFETIQALDVVMRHLPSMRYTPVGRSFFTASEG-----

AGO1_PANTRO     RMLHEALV--------SGQIPVPLESVQALDVAMRHLASMRYTPVGRSFFSPPEG-----
AGO1_MUSMUS     RMLHEALV--------SGQIPVPLESVQALDVAMRHLASMRYTPVGRSFFSPPEG-----
AGO1_XENLA      RMLHEALG--------SGRIQLPLESVQALDVAMRHLASMRYTPVGRSFFSPPEG-----
AGO1_DROME      FNLEEALEG--------RTRQIPYDAILALDVVMRHLPSMTYTPVGRSFFSSPEG-----
AGO1_DANRER     RLLQETLV--------SGRLQVPLGSVQALDVAMRHLASMRYTPVGRSFFSPPEG-----
ALG-1_CAEEL     STLEDAMEG--------RVRQVPFEAVQAMDVILRHLPSLKYTPVGRSFFSPPVPNASGV
AGO1_ARATH      HHLGMFLEG--------KQSDAPQEALQVLDIVLRELPTSRYIPVGRSFYSPDIG-----
AGO1_SACPOM     HTLSQFVNS------KYSSDPQVLSSIMFLDLLLKKKPSETLFGFMHSFFTGENG-----
                  *   :                  .:  :*:  :. .:     . :**::


AGO1            ------------------------YYHPLGGGREVWFGFHQSVRPAMWKMMLNIDVSATA 221
AGO3            ------------------------YDHPLGGGREVWFGFHQSVRPAMWKMMLNIDVSATA
AGO4            ------------------------YYHPLGGGREVWFGFHQSVRPAMWNMMLNIDVSATA
AGO2            ------------------------CSNPLGGGREVWFGFHQSVRPSLWKMMLNIDVSATA

AGO1_PANTRO     ------------------------YYHPLGGGREVWFGFHQSVRPAMWKMMLNIDVSATA
AGO1_MUSMUS     ------------------------YYHPLGGGREVWFGFHQSVRPAMWKMMLNIDVSATA
AGO1_XENLA      ------------------------YYHPLGGGREVWFGFHQSVRPAMWNMMLNIDVSATA
AGO1_DROME      ------------------------YYHPLGGGREVWFGFHQSVRPSQWKMMLNIDVSATA
AGO1_DANRER     ------------------------YYHPLGGGREVWFGFHQSVRPAMWKMMLNIDVSATA
ALG-1_CAEEL     MAGSCPPQASGAVAGGAHSAGQYHAESKLGGGREVWFGFHQSVRPSQWKMMLNIDVSATA
AGO1_ARATH      ------------------------KKQSLGDGLESWRGFYQSIRPTQMGLSLNIDMSSTA
AGO1_SACPOM     --------------------------VSLGGGVEAWKGFYQSIRPNQGFMSVNVDISSSA
                                            **.* * * **:**:**    : :*:*:*::*


AGO1            FYKAQPVIEFMCEVLDIRNIDEQPK-PLTDSQRVRFTKEIKGLKVEVTHCGQMKRKYRVC 280
AGO3            FYKAQPVIQFMCEVLDIHNIDEQPR-PLTDSHRVKFTKEIKGLKVEVTHCGTMRRKYRVC
AGO4            FYRAQPIIEFMCEVLDIQNINEQTK-PLTDSQRVKFTKEIRGLKVEVTHCGQMKRKYRVC
AGO2            FYKAQPVIEFVCEVLDFKSIEEQQK-PLTDSQRVKFTKEIKGLKVEITHCGQMKRKYRVC

AGO1_PANTRO     FYKAQPVIEFMCEVLDIRNIDEQPK-PLTDSQRVRFTKEIKGLKVEVTHCGQMKRKYRVC
AGO1_MUSMUS     FYKAQPVIEFMCEVLDIRNIDEQPK-PLTDSQRVRFTKEIKGLKVEVTHCGQMKRKYRVC
AGO1_XENLA      FYKAQPVIEFMCEVLDIRNIDEQPK-PLTDSQRVRFTKEIKGLKVEVTHCGQMKRKYRVC
AGO1_DROME      FYKAQPVIDFMCEVLDIRDINEQRK-PLTDSQRVKFTKEIKGLKIEITHCGQMRRKYRVC
AGO1_DANRER     FYKAQPVIEFMCEVLDIRNIDEQPK-TLTDSQRVRFTKEIKGLKVEVTHCGQMKRKYRVC
ALG-1_CAEEL     FYRSMPVIEFIAEVLELPVQALAERRALSDAQRVKFTKEIRGLKIEITHCGQMRRKYRVC
AGO1_ARATH      FIEANPVIQFVCDLLNRDISS----RPLSDADRVKIKKALRGVKVEVTHRGNMRRKYRIS
AGO1_SACPOM     FWRNDSLLQILMEYTDCSNVRDLTRFDLKRLSRKFRFLKVTCQHRNNVGTDLANRVYSIE
                * .  .:::: :  :           *.   *      :   : : .  .  .* * :
```

```
AGO1            NVTRRPASHQTFPLQLESGQTVECTVAQYFKQKYNLQLKYPHLPCLQVGQEQKHTYLPLE 340
AGO3            NVTRRPASHQTFPLQLENGQTVERTVAQYFREKYTLQLKYPHLPCLQVGQEQKHTYLPLE
AGO4            NVTRRPASHQTFPLQLENGQAMECTVAQYFKQKYSLQLKYPHLPCLQVGQEQKHTYLPLE
AGO2            NVTRRPASHQTFPLQQESGQTVECTVAQYFKDRHKLVLRYPHLPCLQVGQEQKHTYLPLE

AGO1_PANTRO     NVTRRPASHQTFPLQLESGQTVECTVAQYFKQKYNLQLKYPHLPCLQVGQEQKHTYLPLE
AGO1_MUSMUS     NVTRRPASHQTFPLQLESGQTVECTVAQYFKQKYNLQLKYPHLPCLQVGQEQKHTYLPLE
AGO1_XENLA      NVTRRPASHQTFPLQLESGQTVECTVAQYFKQKYNLQLKYPHLPCLQVGQEQKHTYLPLE
AGO1_DROME      NVTRRPAQMQSFPLQLENGQTVECTVAKYFLDKYRMKLRYPHLPCLQVGQEHKHTYLPLE
AGO1_DANRER     NVTRRPASHQTFPLQLESGQTVECTVAQYFKQKYNLQLKYPHLPCLQVGQEQKHTYLPLE
ALG-1_CAEEL     NVTRRPAQTQTFPLQLETGQTIECTVAKYFYDKYRIQLKYPHLPCLQVGQEQKHTYLPPE
AGO1_ARATH      GLTAVATRELTFPVDERN---TQKSVVEYFHETYGFRIQHTQLPCLQVGNSNRPNYLPME
AGO1_SACPOM     GFSSK-SASDSFFVRRLNGEEQKISVAEYFLENHNVRLQYPNLPCILVKNG---AMLPIE
                ..:   :   :* :   .    : :*.:** : : . :::.:***: * :      ** *


AGO1            VCNIVAGQRCIKKLTDNQTSTMIKATARSAPDRQEEISR-----LMKN--ASYNLDPYIQ 393
AGO3            VCNIVAGQRCIKKLTDNQTSTMIKATARSAPDRQEEISR-----LVR--SANYETDPFVQ
AGO4            VCNIVAGQRCIKKLTDNQTSTMIKATARSAPDRQEEISR-----LVKSNSMVGGPDPYLK
AGO2            VCNIVAGQRCIKKLTDNQTSTMIRATARSAPDRQEEISK-----LMRS--ASFNTDPYVR

AGO1_PANTRO     VCNIVAGQRCIKKLTDNQTSTMIKATARSAPDRQEEISR-----LMKN--ASYNLDPYIQ
AGO1_MUSMUS     VCNIVAGQRCIKKLTDNQTSTMIKATARSAPDRQEEISR-----LMKN--ASYNLDPYIQ
AGO1_XENLA      VCNIVPGQRCIKKLTDNQTSTMIKATARSAPDRQEEISR-----LMKN--ASYNLDPYIQ
AGO1_DROME      VCNIVAGQRCIKKLTDMQTSTMIKATARSAPDREREINN-----LVKR--ADFNNDSYVQ
AGO1_DANRER     VCNIVAGQRCIKKLTDNQTSTMIKATARSAPDRQEEISR-----LMKN--ANFNLDPYIQ
ALG-1_CAEEL     VCNIVPGQRCIKKLTDVQTSTMIKATARSAPERERISN-----LVRK--AEFSADPFAH
AGO1_ARATH      VCKIVEGQRYSKRLNERQITALLKVTCQRPIDREKDILQTVQLN-------DYAKDNYAQ
AGO1_SACPOM     FCFVVKGQRYTAKLNSDQTANMIRFAVQRPFERVQQIDD-------FVHQMDWDTDPYLT
                .* :* ***   :*.. * : ::: : : . :* .:*                  * :


AGO1            EFGIKVKDDMTEVTGRVLPAPILQYGGRN---------------RAIATPNQGVWDMRGK 438
AGO3            EFQFKVRDEMAHVTGRVLPAPMLQYGGRN---------------RTVATPSHGVWDMRGK
AGO4            EFGIVVHNEMTELTGRVLPAPMLQYGGRN---------------KTVATPNQGVWDMRGK
AGO2            EFGIMVKDEMTDVTGRVLQPPSILYGGRN---------------KAIATPVQGVWDMRNK

AGO1_PANTRO     EFGIKVKDDMTEVTGRVLPAPILQYGGRN---------------RAIATPNQGVWDMRGK
AGO1_MUSMUS     EFGIKVKDDMTEVTGRVLPAPILQYGGRN---------------RAIATPNQGVWDMRGK
AGO1_XENLA      EFGIKVKDDMTEVTGRVLPAPILQYGGRN---------------RAIATPNQGVWDMRGK
AGO1_DROME      EFGLTISNSMMEVRGRVLPPPKLQYGGRVSTGLTGQQLFPPQNKVSLASPNQGVWDMRGK
AGO1_DANRER     EFGIKVKDDMTEVTGRVLPAPILQYGGRN---------------RAIATPNQGVWDMRGK
ALG-1_CAEEL     EFGITINPAMTEVKGRVLSAPKLLYGGRT---------------RATALPNQGVWDMRGK
AGO1_ARATH      EFGIKISTSLASVEARILPPPWLKYHESG--------------REGTCLPQVGQWNMMNK
AGO1_SACPOM     QYGMKIQKKMLEVPARVLETPSIRYGGDC-----------------IERPVSGRWNLRGK
                :: : :   :  : .*:* .* : *                        *  * *:: .*


AGO1            QFYNG--IEIKVWAIACFAPQKQCREEVLKNFTDQLRKISKDAGMPIQGQPCFCKYAQGA 496
AGO3            QFHTG--VEIKMWAIACFATQRQCREEILKGFTDQLRKISKDAGMPIQGQPCFCKYAQGA
AGO4            QFYAG--IEIKVWAVACFAPQKQCREDLLKSFTDQLRKISKDAGMPIQGQPCFCKYAQGA
AGO2            QFHTG--IEIKVWAIACFAPQRQCTEVHLKSFTEQLRKISRDAGMPIQGQPCFCKYAQGA

AGO1_PANTRO     QFYNG--IEIKVWAIACFAPQKQCREEVLKNFTDQLRKISKDAGMPIQGQPCFCKYAQGA
AGO1_MUSMUS     QFYNG--IEIKVWAIACFAPQKQCREEVLKNFTDQLRKISKDAGMPIQGQPCFCKYAQGA
AGO1_XENLA      QFYNG--IEIKVWAIACFAPQKQCREEVLKNFTDQLRKISKDAGMPIQGQPCFCKYAQGA
AGO1_DROME      QFFTG--VEIRIWAIACFAPQRTVREDALRNFTQQLQKISNDAGMPIIGQPCFCKYATGP
AGO1_DANRER     QFYNG--IEIKVWAIACFAPQKQCREEVLKNFTDQLRKISKDAGMPIQGQPCFCKYAQGA
ALG-1_CAEEL     QFHTG--IDVRVWAIACFAQQQHVKENDLRMFTNQLQRISNDAGMPIVGNPCFCKYAVGV
AGO1_ARATH      KMING--GTVNNWICINFS--RQVQDNLARTFCQELAQMCYVSGMAFNPEPVLPPVSARP
AGO1_SACPOM     RFLDPPRAPIRSWAVMCFTSTRRLPMRGIENFLQTYVQTLTSLGINFVMKKPPVLYADIR
                ::       :. *    *:  :       . * :   :     *: :  :      :
```

```
AGO1            DSVEPMFRHLKNTYSG--------LQLIIVILPGKTPVYAEVKRVGDTLLGMATQCVQVK 548
AGO3            DSVEPMFRHLKNTYSG--------LQLIIVILPGKTPVYAEVKRVGDTLLGMATQCVQVK
AGO4            DSVEPMFKHLKMTYVG--------LQLIVVILPGKTPVYAEVKRVGDTLLGMATQCVQVK
AGO2            DSVEPMFRHLKNTYAG--------LQLVVVILPGKTPVYAEVKRVGDTVLGMATQCVQMK

AGO1_PANTRO     DSVEPMFRHLKNTYSG--------LQLIIVILPGKTPVYAEVKRVGDTLLGMATQCVQVK
AGO1_MUSMUS     DSVEPMFRHLKNTYSG--------LQLIIVILPGKTPVYAEVKRVGDTLLGMATQCVQVK
AGO1_XENLA      DSVEPMFRHLKNTYSG--------LQLIIVILPGKTPVYAEVKRVGDTLLGMATQCVQVK
AGO1_DROME      DQVEPMFRYLKITFPG--------LQLVVVVLPGKTPVYAEVKRVGDTVLGMATQCVQAK
AGO1_DANRER     DSVEPMFRHLKNTYSG--------LQLIIVILPGKTPVYAEVKRVGDTLLGMATQCVQVK
ALG-1_CAEEL     EQVEPMFKYLKQNYSG--------IQLVVVVLPGKTPVYAEVKRVGDTVLGIATQCVQAK
AGO1_ARATH      EQVEKVLKTRYHDATSKLSQGKEIDLLIVILPDNNGSLYGDLKRICETELGIVSQCCLTK
AGO1_SACPOM     GSVEELCITLYKKAEQVG--NAPPDYLFFILDKNSPEPYGSIKRVCNTMLGVPSQCAISK
                 .** :                    *..::  ..   *..:**: :* **: :**   *


AGO1            NVVKTSPQTLSNLCLKINVKLGGINNILVPHQR---SAVFQQPVIFLGADVTHPPAGDGK 605
AGO3            NVIKTSPQTLSNLCLKINVKLGGINNILVPHQR---PSVFQQPVIFLGADVTHPPAGDGK
AGO4            NVVKTSPQTLSNLCLKINAKLGGINNVLVPHQR---PSVFQQPVIFLGADVTHPPAGDGK
AGO2            NVQRTTPQTLSNLCLKINVKLGGVNNILLPQGR---PPVFQQPVIFLGADVTHPPAGDGK

AGO1_PANTRO     NVVKTSPQTLSNLCLKINVKLGGINNILVPHQR---SAVFQQPVIFLGADVTHPPAGDGK
AGO1_MUSMUS     NVVKTSPQTLSNLCLKINVKLGGINNILVPHQR---SAVFQQPVIFLGADVTHPPAGDGK
AGO1_XENLA      NVVKTSPQTLSNLCLKINVKLGGINNILVPHQR---SAVFQQPVIFLGADVTHPPAGDGK
AGO1_DROME      NVNKTSPQTLSNLCLKINVKLGGINSILVPSIR---PKVFNEPVIFLGADVTHPPAGDNK
AGO1_DANRER     NVVKTSPQTLSNLCLKINVKLGGINNILVPHQR---SAVFQQPVIFLGADVTHPPAGDGK
ALG-1_CAEEL     NAIRTTPQTLSNLCLKMNVKLGGVNSILLPNVR---PRIFNEPVIFFGCDITHPPAGDSR
AGO1_ARATH      HVFKMSKQYMANVALKINVKVGGRNTVLVDALSRRIPLVSDRPTIIFGADVTHPHPGEDS
AGO1_SACPOM     HILQSKPQYCANLGMKINVKVGGINCSLIPKSN----PLGNVPTLILGGDVYHPGVG-AT
                :  : . *  :*: :*:*.*:** *  *:         : : *.:::* *: **  *


AGO1            KPSITAVVGSMDAHPS-RYCATVRVQRPRQ-------------EIIEDLSYMVRELLIQF 651
AGO3            KPSIAAVVGSMDAHPS-RYCATVRVQRPRQ-------------EIIQDLASMVRELLIQF
AGO4            KPSIAAVVGSMDGHPS-RYCATVRVQTSRQEISQELLYSQ---EVIQDLTNMVRELLIQF
AGO2            KPSIAAVVGSMDAHPN-RYCATVRVQQHRQ-------------EIIQDLAAMVRELLIQF

AGO1_PANTRO     KPSITAVVGSMDAHPS-RYCATVRVQRPRQ-------------EIIEDLSYMVRELLIQF
AGO1_MUSMUS     KPSITAVVGSMDAHPS-RYCATVRVQRPRQ-------------EIIEDLSYMVRELLIQF
AGO1_XENLA      KPSITAVVGSMDAHPS-RYCATVRVQRPRQ-------------EIIEDLSYMVRELLIQF
AGO1_DROME      KPSIAAVVGSMDAHPS-RYAATVRVQQHRQ-------------EIIQELSSMVRELLIMF
AGO1_DANRER     KPSITAVVGSMDAHPS-RYCATVRVQRPRQ-------------EIIEDLSYMVRELLIQF
ALG-1_CAEEL     KPSIAAVVGSMDAHPS-RYAATVRVQQHRQ-------------EIISDLTYMVRELLVQF
AGO1_ARATH      SPSIAAVVASQDWPEITKYAGLVCAQAHRQELIQDLFKEWKDPQKGVVTGGMIKELLIAF
AGO1_SACPOM     GVSIASIVASVDLNGC-KYTAVSRSQPRHQ-------------EVIEGMKDIVVYLLQGF
                  **:::*.* *     :* .    *  :*             :       ::  **   *


AGO1            YKSTR-FKPTRIIFYRDGVPEGQLPQILHYELLAIRDACIKLEKDYQPGITYIVVQKRHH 710
AGO3            YKSTR-FKPTRIIFYRDGVSEGQFRQVLYYELLAIREACISLEKDYQPGITYIVVQKRHH
AGO4            YKSTR-FKPTRIIYYRGGVSEGQMKQVAWPELIAIRKACISLEEDYRPGITYIVVQKRHH
AGO2            YKSTR-FKPTRIIFYRDGVSEGQFQQVLHHELLAIREACIKLEKDYQPGITFIVVQKRHH

AGO1_PANTRO     YKSTR-FKPTRIIFYRDGVPEGQLPQILHYELLAIRDACIKLEKDYQPGITYIVVQKRHH
AGO1_MUSMUS     YKSTR-FKPTRIIFYRDGVPEGQLPQILHYELLAIRDACIKLEKDYQPGITYIVVQKRHH
AGO1_XENLA      YKSTR-FKPTRIIFYRDGVPEGQLPQILHYELLAIRDACIKLEKDYQPGITYIVVQKRHH
AGO1_DROME      YKSTGGYKPHRIILYRDGVSEGQFPHVLQHELTAIREACIKLEPEYRPGITFIVVQKRHH
AGO1_DANRER     YKSTR-FKPTRIIFYRDGVPEGQLPQILHYELLAIRDACIKLEKDYQPGITYIVVQKRHH
ALG-1_CAEEL     YRNTR-FKPARIVVYRDGVSEGQFFNVLQYELRAIREACMMLERGYQPGITFIAVQKRHH
AGO1_ARATH      RRSTG-HKPLRIIFYRDGVSEGQFYQVLLYELDAIRKACASLEAGYQPPVTFVVVQKRHH
AGO1_SACPOM     RAMTK-QQPQRIIYFRDGTSEGQFLSVINDELSQIKEACHSLSPKYNPKILVCTTQKRHH
                   *   :* **: :*.*..***:  :   **  *:.**  *.  *.* :   ..*****
```

```
AGO1            TRLFCADKNE--RIGKSGNIPAGTTVDTNITHPFEFDFYLCSHAGIQGTSRPSHYYVLWD 768
AGO3            TRLFCADRTE--RVGRSGNIPAGTTVDTDITHPYEFDFYLCSHAGIQGTSRPSHYHVLWD
AGO4            TRLFCADKTE--RVGKSGNVPAGTTVDSTITHPSEFDFYLCSHAGIQGTSRPSHYQVLWD
AGO2            TRLFCTDKNE--RVGKSGNIPAGTTVDTKITHPTEFDFYLCSHAGIQGTSRPSHYHVLWD

AGO1_PANTRO     TRLFCADKNE--RIGKSGNIPAGTTVDTNITHPFEFDFYLCSHAGIQGTSRPSHYYVLWD
AGO1_MUSMUS     TRLFCADKNE--RIGKSGNIPAGTTVDTNITHPFEFDFYLCSHAGIQGTSRPSHYYVLWD
AGO1_XENLA      TRLFCADRSE--RIGKSGNIPAGTTVDTNITHPFEFDFYLCSHAGIQGTSRPSHYYVLWD
AGO1_DROME      TRLFCAEKKE--QSGKSGNIPAGTTVDVGITHPTEFDFYLCSHQGIQGTSRPSHYHVLWD
AGO1_DANRER     TRLFCADKSE--RIGKSGNIPAGTTVDTSITHPFEFDFYLCSHAGIQGTSRPSHYYVLWD
ALG-1_CAEEL     TRLFAVDKKD--QVGKAYNIPPGTTVDVGITHPTEFDFYLCSHAGIQGTSRPSHYHVLWD
AGO1_ARATH      TRLFAQNHNDRHSVDRSGNILPGTVVDSKICHPTEFDFYLCSHAGIQGTSRPAHYHVLWD
AGO1_SACPOM     ARFFIKNKSD---GDRNGNPLPGTIIEKHVTHPYQYDFYLISHPSLQGVSVPVHYTVLHD
                :*:*  ::.:    .:  *  .** ::  : ** ::**** ** .:**.* * ** ** *


AGO1            DNRFTADELQILTYQLCHTYVRCTRSVSIPAPAYYARLVAFRARYHLVDKEHDSG----- 823
AGO3            DNCFTADELQLLTYQLCHTYVRCTRSVSIPAPAYYAHLVAFRARYHLVDKEHDSA-----
AGO4            DNCFTADELQLLTYQLCHTYVRCTRSVSIPAPAYYARLVAFRARYHLVDKDHDSA-----
AGO2            DNRFSSDELQILTYQLCHTYVRCTRSVSIPAPAYYAHLVAFRARYHLVDKEHDSA-----

AGO1_PANTRO     DNRFTADELQILTYQLCHTYVRCTRSVSIPAPAYYARLVAFRARYHLVDKEHDSG-----
AGO1_MUSMUS     DNRFTADELQILTYQLCHTYVRCTRSVSIPAPAYYARLVAFRARYHLVDKEHDSG-----
AGO1_XENLA      DNRFTADELQILTYQLCHTYVRCTRSVSIPAPAYYARLVAFRARYHLVDKEHDSG-----
AGO1_DROME      DNHFDSDELQCLTYQLCHTYVRCTRSVSIPAPAYYAHLVAFRARYHLVEKEHDSG-----
AGO1_DANRER     DNRFTADELQILTYQLCHTYVRCTRSVSIPAPAYYARLVAFRARYHLVDKEHDSG-----
ALG-1_CAEEL     DNNLTADELQQLTYQMCHTYVRCTRSVSIPAPAYYAHLVAFRARYHLVDREHDSG-----
AGO1_ARATH      ENNFTADGLQSLTNNLCYTYARCTRSVSIVPPAYYAHLAAFRARFYMEPETSDSGSMASG
AGO1_SACPOM     EIQMPPDQFQTLCYNLCYVYARATSAVSLVPPVYYAHLVSNLARYQDVTADDTFV-----
                :  : .* :* *  ::*:.*.*.* :**: .*.***:*.:  **:


AGO1            ----EGSHISGQSNGRDPQALAKAVQVHQDTLRTMYFA- 857
AGO3            ----EGSHVSGQSNGRDPQALAKAVQIHQDTLRTMYFA-
AGO4            ----EGSHVSGQSNGRDPQALAKAVQIHHDTQHTMYFA-
AGO2            ----EGSHTSGQSNGRDHQALAKAVQVHQDTLRTMYFA-

AGO1_PANTRO     ----EGSHISGQSNGRDPQALAKAVQVHQDTLRTMYFA-
AGO1_MUSMUS     ----EGSHISGQSNGRDPQALAKAVQVHQDTLRTMYFA-
AGO1_XENLA      ----EGSHISGQSNGRDPQALAKAVQVHQDTLRTMYFA-
AGO1_DROME      ----EGSHQSGCSEDRTPGAMARAITVHADTKKVMYFA-
AGO1_DANRER     ----EGSHVSGQSNGRDPQALAKAVQIHHDSLRTMYFA-
ALG-1_CAEEL     ----EGSQPSGTSEDTTLSNMARAVQVHPDANNVMYFA-
AGO1_ARATH      SMARGGGMAGRSTRGPNVNAAVRPLPALKENVKRVMFYC
AGO1_SACPOM     -----------ETSEASMDQEVKPLLALSSKLKTKMWYM
                            :        .:.:    .  .   :
```

### Figure S3

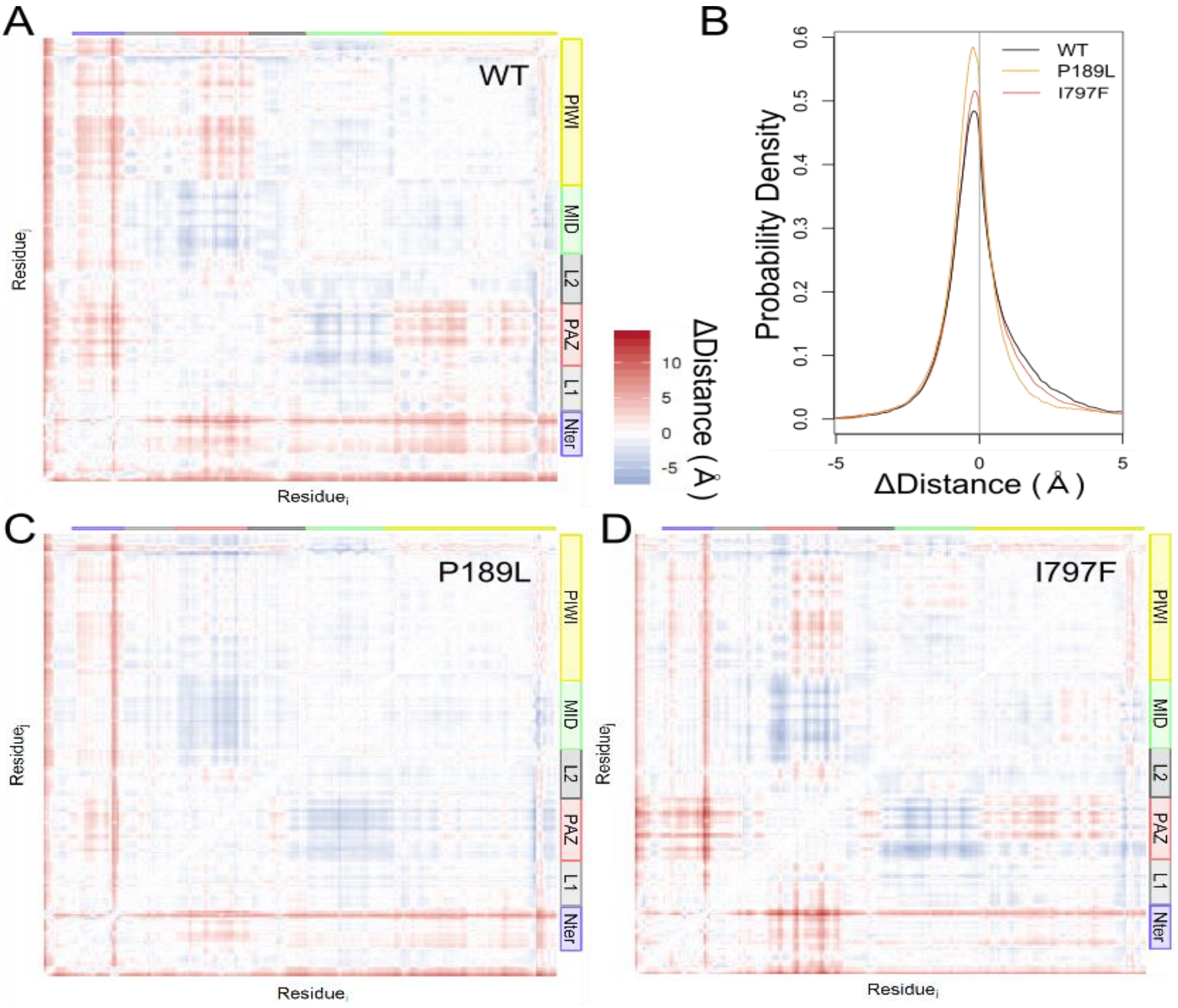
A
WT
PIWI
MID
L2
PAZ
L1
Nter
Residue_j
Residue_i
ΔDistance ( Å )
10
5
0
-5
B
WT
P189L
I797F
Probability Density
0.0
0.1
0.2
0.3
0.4
0.5
0.6
-5
0
5
ΔDistance ( Å )
C
P189L
D
I797F

**Figure S4**

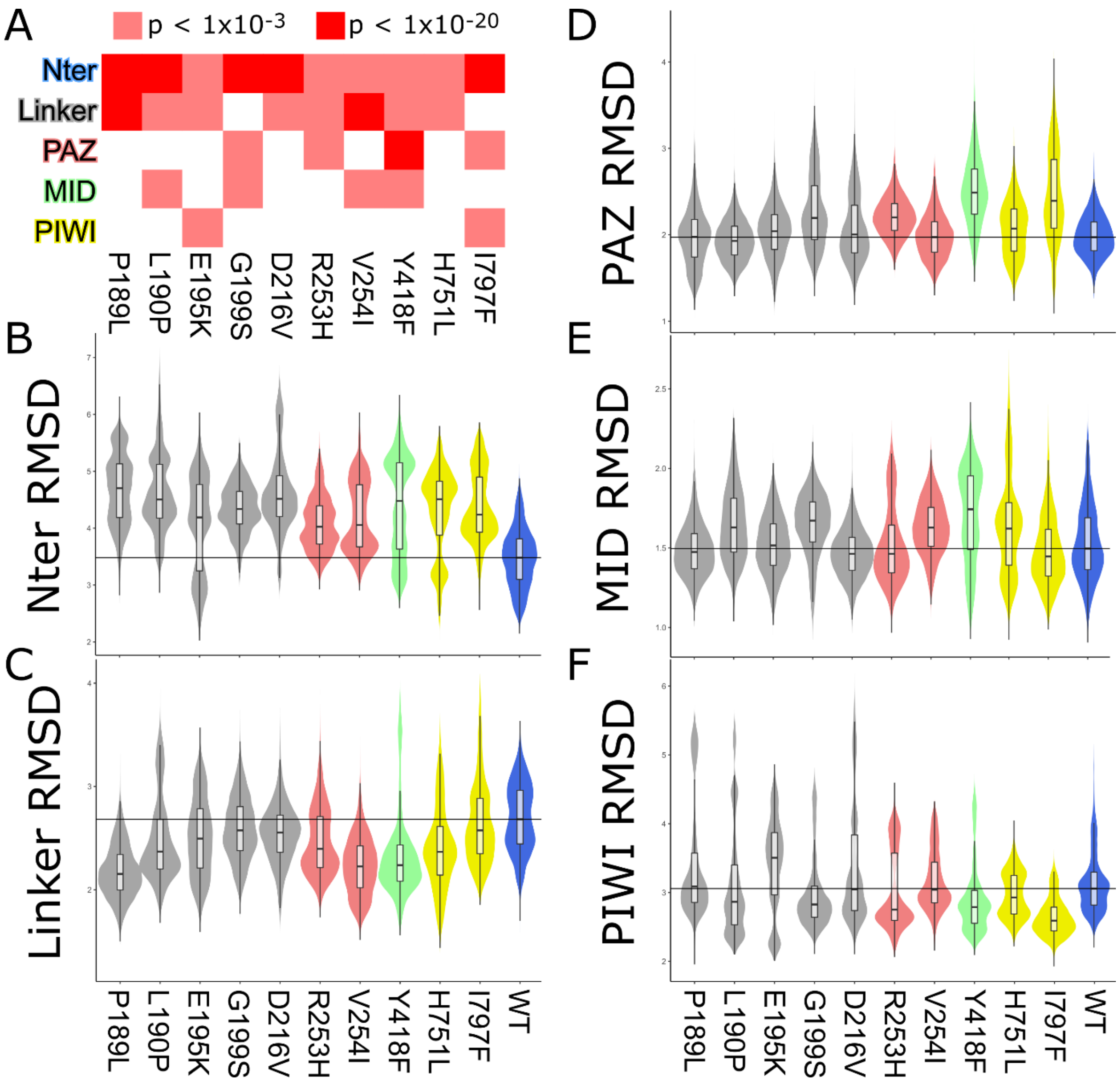

A
p < 1x10-3
p < 1x10-20
Nter
Linker
PAZ
MID
PIWI
P189L
L190P
E195K
G199S
D216V
R253H
V254I
Y418F
H751L
I797F
B
Nter RMSD
C
Linker RMSD
D
PAZ RMSD
E
MID RMSD
F
PIWI RMSD
WT

| | Family 1 | Family 2 | Family 3 | Family 4 | Family 5 | Family 6 | Family 7 | Family 8 | Rauch *et al.*[13] |
|---|---|---|---|---|---|---|---|---|---|
| **NM_012199.4 (*AGO1*)** | c.539_541delTCT p.(Phe180del) | c.539_541delTCT p.(Phe180del) | c.539_541delTCT p.(Phe180del) | c.539_541delTCT p.(Phe180del) | c.539_541delTCT p.(Phe180del) | c.566C>T p.(Pro189Leu) | c.569T>G p.(Leu190Arg) | c.569T>G p.(Leu190Arg) | c.569T>C p.(Leu190Pro) |
| **Variation status** | *de novo* | *de novo* | *de novo* | *de novo* | *de novo* | *de novo* | *de novo* | *de novo* | ND |
| **Protein domain** | L1 | L1 | L1 | L1 | L1 | L1 | L1 | L1 | L1 |
| **Current age** (years) | 10.8 | 3 | 8 | 8 | 6 | 9 | 8.8 | 12 | ND |
| **Gender** | M | M | M | M | F | M | F | F | F |
| Birth | | | | | | | | | |
| **Age of birth** (weeks) | 41 | 37 | 38 | 37.1 | ND | 41 | 35 (twin dz) | 41 | 42 |
| **Weight** | - 0.3 SD | - 1.8 SD | - 0.5 SD | - 2.3 SD | - 1.6 SD | + 0.4 SD | - 1.2 SD | ND | N |
| **Length** | - 0.3 SD | ND | - 1 SD | - 1.18 SD | ND | ND | - 1.8 SD | + 0.2 SD | ND |
| **OFC** | ND | ND | - 0.2 SD | - 1.71 SD | ND | ND | - 0.2 SD | ND | ND |
| Growth | | | | | | | | | |
| **Weight** | + 0 SD | + 0.3 SD at 2 yrs | - 2.7 SD at 7 yrs | - 2.33 SD | - 3.9 SD | + 2 SD | - 3 SD | - 0.2 SD | ND |
| **Length** | + 0.6 SD | 0 SD at 2 yrs | - 2.8 SD at 7 yrs | - 2.77 SD | - 3.3 SD | ND | - 2.7 SD | - 3.1 SD | -1.8 SD |
| **OFC** | + 0.6 SD | + 1 SD at 2 years | + 0.5 SD at 7 yrs | - 1 SD | - 0.6 SD | + 0.3 SD | - 1.5 SD | + 1.2 SD | -0.9 SD |
| Clinical characteristics | | | | | | | | | |
| **ID/DD** | mild | moderate | severe | + | ND | moderate | severe | moderate | severe |
| **Hypotonia** | ND | + | ND | ND | + | ND | ND | + | + |
| **History of seizures** | + | - | + | - | - | ND | + | + | - |
| **Motor delay** | + | + | + | + | + | + | + | + | + |
| Walk (months) | ND | 24 | 20 | 18 | no at 24 | ND | 36-40 | 24 | 48 |
| **Speech delay** | + | + | + | + | + | +[b] | + | + | + |
| 1st words (months) | 12-24 | no at 24 | ND | ND | ND | no (regression) | no | 42 | ND |
| **Autistic behaviour** | + | + | + | + | + | + | + | + | ND |
| Stereotypies | +[c] | ND | ND | + | +[c] | + | ND | ND | ND |
| **Behaviour disorders** | | | | | | | | | |
| Hyperactivity | + | ND | - | + | - | + | ND | + | ND |
| Anxiety | + | ND | ND | ND | ND | ND | ND | ND | ND |
| Aggressiveness | + | ND | ND | + | ND | + | + | + | ND |
| **Feeding difficulties** | - | ND | + | + | ND | - | - | - | ND |
| **Sleeping disturbance** | + | ND | + | + | + | + | - | + | ND |
| **Facial findings** | | | | | | | | | |
| Prominent forehead | - | - | - | ND | ND | - | + | ND | ND |
| almond eyes | - | - | + | ND | ND | - | - | + | ND |
| flat nasal bridge | - | - | + | ND | ND | - | - | ND | ND |
| Thin upper lip | - | - | + | ND | ND | - | - | ND | ND |
| Other | no | no | epicanthus | wide mouth, high | Small nose | no | telecanthus; | hyperlordosis | ND |
| **Other symptoms** | hypothyroidy, growth horm defic | plagiocephaly, brachycephaly | hypothyroidy, periodic fever | MRI: CCH, bilateral hippocampal dysplasia | hypoplastic teeth, reduced enamel | fever at 18month | acetab dysplasia | supernum. nipple, strabismus; MRI: CCA, colpocephaly | ND |

| | Family 9 | | | Family 10 | Family 11 | Family 12 | Family 13 | Family 14 |
|---|---|---|---|---|---|---|---|---|
| | mz twin 1 | mz twin 2 | Martinez *et al.*[14] | | | Hamdan *et al.*[9] | | |
| **NM_012199.4 (*AGO1*)** | c.569T>C p.(Leu190Pro) | c.569T>C p.(Leu190Pro) | c.583G>A p.(Glu195Lys) | c.595G>A p.(Gly199Ser) | c.595G>A p.(Gly199Ser) | c.595G>A p.(Gly199Ser) | c.595G>A p.(Gly199Ser) | c.595G>A p.(Gly199Ser) |
| **Variation status** | *de novo* | *de novo* | ND | *de novo* | *de novo* | *de novo* | *de novo* | *de novo* |
| **Protein domain** | L1 | L1 | L1 | L1 | L1 | L1 | L1 | L1 |
| **Current age** (years) | 8 | 8 | ND | 8 | 10.5 | 12.5 | 10 | 5.6 |
| **Gender** | F | F | M | F | F | F | F | M |
| Birth | | | | | | | | |
| **Age of birth** (weeks) | 37+6 (twin mz) | 37+6 (twin mz) | ND | 39 | 41 | ND | 42 | 41 |
| **Weight** | - 1 SD | - 1 SD | ND | 0 SD | + 0.8 SD | ND | + 2.9 SD | + 0.8 SD |
| **Length** | - 1 SD | - 1,5 SD | ND | ND | ND | ND | + 2.2 SD | + 2 SD |
| **OFC** | + 0.5 SD | + 0.5 SD | ND | ND | ND | ND | + 3.3 SD | + 0.7 SD |
| Growth | | | | | | | | |
| **Weight** | - 1,5 SD | - 2 SD | increase | 0 SD at 2 years | - 1.2 SD | + 2.5 SD | - 0.9 SD | + 4 |
| **Length** | - 3 SD | - 3 SD | tall stature | - 0.5 SD at 7 years | - 3.1 SD | - 0.4 SD | - 1.6 SD | + 1.5 SD |
| **OFC** | + 1 SD | + 0 SD | ND | + 1.8 SD at 8 years | + 0.6 SD | + 0.1 SD | + 0.2 SD | - 0.6 SD |
| Clinical characteristics | | | | | | | | |
| **ID/DD** | severe | severe | mild | mild | moderate | moderate | moderate | moderate |
| **Hypotonia** | + | + | + | ND | + | ND | + | - |
| **History of seizures** | + | + | - | - | +[a] | +[a] | + | - |
| **Motor delay** | + | + | + | + | + | + | + | + |
| Walk (months) | ND | ND | ND | 20 | 20 | 24 | 36 | 20 |
| **Speech delay** | + | + | + | + | + | + | + | + |
| 1st words (months) | ND | ND | ND | 42 | 12 | 36 | 18 | No at 36 |
| **Autistic behaviour** | + | + | + | - | + | - | - | + |
| Stereotypies | + | + | + | ND | ND | ND | - | ND |
| **Behaviour disorders** | | | | | | | | |
| Hyperactivity | ND | ND | + | - | - | ND | - | + |
| Anxiety | ND | ND | ND | ND | ND | ND | - | ND |
| Aggressiveness | ND | ND | ND | ND | ND | ND | - | ND |
| **Feeding difficulties** | + | + | ND | + | - | ND | - | - |
| **Sleeping disturbance** | ND | ND | + | + | - | ND | + | + |
| **Facial findings** | | | | | | | | |
| Prominent forehead | + | + | ND | + | ND | ND | + | + |
| Almond eyes | + | + | ND | + | ND | ND | + | + |
| flat nasal bridge | - | - | + | - | ND | ND | - | + |
| Thin upper lip | + | + | + | + | ND | ND | + | + |
| Other | no | no | large ears, | no | no | no | no | no |
| **Other symptoms** | profound deafness, absent eye contact | mild deafness, absent eye contact | kidney abnormality | no | no | no | no | brachycephaly |

| | Sakaguchi *et al.*[15] | DDD Study[16] | **Family 15** | **Family 16** | **Family 17** | **Family 18** | **Family 19** | Sanders *et al.*[17] | **Family 20** |
|---|---|---|---|---|---|---|---|---|---|
| **NM_012199.4 (*AGO1*)** | c.595G>A p.(Gly199Ser) | c.647A>T p.(Asp216Val) | c.650-2A>G p.(Val217_Ser218del) | c.758G>A p.(Arg253His) | c.760G>A p.(Val254Ile) | c.760G>A p.(Val254Ile) | c.971C>T p.(Pro324Leu) | c.1064C>T p.(Thr355Ile) | c.1073A>G p.(Gln358Arg) |
| **Variation status** | *de novo* | ND | *de novo* | *de novo* | *de novo* | *de novo* | *de novo* | ND | *de novo* |
| **Protein domain** | L1 | L1 | | PAZ | PAZ | PAZ | PAZ | PAZ | PAZ |
| **Current age** (years) | 15 | ND | 12 | 20 | 15 | 6.5 | 2.3 | ND | 4 |
| **Gender** | F | F | M | M | M | F | M | M | M |
| Birth | | | | | | | | | |
| **Gestation at birth** | 41 | ND | 40 | ND | 41 | ND | ND | ND | 39 |
| **Weight** | + 2.4 SD | ND | + 0.5 SD | ND | + 0.2 SD | - 0.2 SD | - 1.5 SD | ND | + 0.5 SD |
| **Length** | + 1.5 SD | ND | - 0.8 SD | ND | - 1 SD | ND | - 0.5 SD | ND | - 0.9 SD |
| **OFC** | + 1.1 SD | ND | - 0.9 SD | ND | + 1.2 SD | ND | - 2 SD | ND | + 0.6 SD |
| Growth | | | | | | | | | |
| **Weight** | − 1.2 SD | ND | - 1 SD | + 4.1 SD | - 0.3 SD | + 0.5 SD | - 1 SD | ND | - 1.8 SD |
| **Length** | − 2.6 SD | ND | - 0.6 SD | + 0.3 SD | - 0.5 SD | - 0.5 SD | - 4.8 SD | ND | - 2.5 SD |
| **OFC** | + 1.0 SD | ND | - 1.8 SD | + 2.3 SD | + 1.9 SD | + 1,3 SD | + 0.5 SD | ND | + 0.1 SD |
| Clinical characteristics | | | | | | | | | |
| **ID/DD** | moderate | + | moderate | moderate | mild | mild | Moderate | ND | severe |
| **Hypotonia** | + | ND | + | ND | - | - | ND | ND | - |
| **History of seizures** | ND | ND | - | + | + | + | - | ND | - |
| **Motor delay** | + | ND | + | ND | - | - | + | ND | + |
| Walk (months) | ND | ND | 22 | ND | 12 | ND | 21 | ND | 30 |
| **Speech impairment** | ND | ND | + | + | + | + | + | ND | + |
| 1st words (months) | ND | ND | 42 | ND | 36 | ND | - | ND | 16 |
| **Autistic behaviour** | ND | ND | - | + | + | - | + | + | + |
| Stereotypies | ND | ND | - | ND | ND | ND | + | ND | ND |
| **Behaviour disorders** | | | | | | | | | |
| Hyperactivity | ND | ND | + | ND | + | + | - | ND | - |
| Anxiety | ND | ND | ND | ND | ND | + | ND | ND | ND |
| Aggressiveness | ND | ND | ND | + | ND | + | + | ND | - |
| **Feeding difficulties** | ND | ND | - | ND | - | - | ND | ND | - |
| **Sleeping disturbance** | ND | ND | - | + | - | - | + | ND | - |
| **Facial findings** | | | | | | | | | |
| Prominent forehead | ND | ND | ND | ND | ND | ND | + | ND | ND |
| Almond eyes | + | ND | ND | ND | ND | ND | ND | ND | ND |
| Flat nasal bridge | + | ND | ND | ND | ND | + | ND | ND | ND |
| Thin upper lip | ND | ND | ND | ND | ND | ND | - | ND | + |
| Other | telecanthus | ND | deep set ears | ND | ND | epicanthus | macrocephaly | ND | ND |
| **Other symptoms** | ND | ND | mild ataxia, abnormality of the kidney | supernum. nipple | fetal pads | no | ND | ND | phimosis, glabellar hémangioma, 2 café au lait spots |

| | Family 21 | Family 22 | Family 23 | Family 24 | | Family 25 | Family 26 |
|---|---|---|---|---|---|---|---|
| | | | | mz twin 1 | mz twin 2 | twin | |
| **NM_012199.4 (*AGO1*)** | c.1126_1128del p.(Glu376del) | c.1126_1128del p.(Glu376del) | c.1253A>T p.(Tyr418Phe) | c.2252A>T p.(His751Leu) | c.2252A>T p.(His751Leu) | c.2342C>T p.(Thr781Met) | c.2389A>T p.(Ile797Phe) |
| **Variation status** | *de novo* | *de novo* | Unknown | *de novo* | *de novo* | Unknown | *de novo* |
| **Protein domain** | L2 | L2 | L2 | Piwi | Piwi | Piwi | Piwi |
| **Current age** (years) | 9 | 3.4 | 9 | 18 | 18 | 7 | 9 |
| **Gender** | M | F | M | F | F | M | M |
| Birth | | | | | | | |
| **Gestation at birth** | 40 | 41 | ND | 34 (twin mz) | 34 (twin mz) | 36 (twin) | 39 |
| **Weight** | +0/13 SD | - 0.2 SD | + 0.4 SD | - 0,5 SD | + 0.3 SD | - 2.2 SD | + 0.9 SD |
| **Length** | ND | - 1.3 SD | ND | - 2,3 SD | - 1.1 SD | ND | + 0.5 SD |
| **OFC** | -1.3 SD | - 3 SD | ND | < - 2 SD | < - 2 SD | ND | + 1 SD |
| Growth | | | | | | | |
| **Weight** | - 1.7 SD | - 1.7 SD | + 5.5 SD | - 2 SD | - 1.8 SD | - 0.9 SD | - 1.35 SD |
| **Length** | - 2.6 SD | - 1.6 SD | + 1.8 SD | - 1.5 SD | - 0.9 SD | - 1.7 SD | - 1.1 SD |
| **OFC** | -2.07 SD | ND | + 1.2 SD | - 3.9 SD | - 3.9 SD | - 1.8 SD | + 1.5 SD |
| Clinical characteristics | | | | | | | |
| **ID/DD** | severe | mild | borderline | severe | moderate | borderline | mild |
| **Hypotonia** | + | + | ND | ND | ND | ND | - |
| **History of seizures** | + | - | ND | - | - | - | - |
| **Motor delay** | + | + | + | + | + | + | + |
| Walk (months) | 35 | 41 | 18 | 25 | 25 | 18 | 18 |
| **Speech impairment** | + | + | + | + | + | + | + |
| 1st words (months) | no | 19 | ND | 36-42 | 36-42 | 36-48 | ND |
| **Autistic behaviour** | + | + | + | + | + | + | - |
| Stereotypies | + | + | ND | ND | ND | + | - |
| **Behaviour disorders** | | | | | | | |
| Hyperactivity | + | ND | + | + | + | + | + |
| Anxiety | ND | ND | + | + | + | + | + |
| Aggressiveness | ND | ND | + | + | ND | + | - |
| **Feeding difficulties** | + | + | - | + | + | + | - |
| **Sleeping disturbance** | + | ND | + | + | + | ND | + |
| **Facial findings** | | | | | | | |
| Prominent forehead | ND | + | ND | + | + | ND | - |
| Almond eyes | + | ND | ND | + | + | ND | - |
| Flat nasal bridge | ND | ND | + | + | + | + | - |
| Thin upper lip | + | ND | ND | + | + | - | - |
| Other | epicanthus, | ND | ND | no | no | epicanthus, | strabismus |
| **Other symptoms** | ND | metatarsus adductus, lymphedema feet | no | camptodactyly | camptodactyly | nystagmus | hypermetropia, gastro-esophageal reflux during first months of life |

#### Table S2

| | Variants in *AGO1* | deletion 1p34.3 | Tokita *et al.,* 2014 | | | | |
|---|---|---|---|---|---|---|---|
| | | | **proband 1** | **proband 2** | **proband 3** | **proband 4** | **proband 5** |
| **Variations (hg19)** | | | arr[GRCh37] 1p34.3 (36358320_39088512)x1 | arr[GRCh37] 1p34.3 (36154687_38591548)x1 | arr[GRCh37] 1p34.3 (35933018_37052682)x1 | arr[GRCh37] 1p34.3 (35933018_37052682)x1 | arr[GRCh37] 1p34.3 (35447244_36643150)x1 |
| **AGO genes included** | | | *AGO1*, *AGO3* | *AGO1*, *AGO3* and *AGO4* | *AGO1*, *AGO3* and *AGO4* | *AGO1*, *AGO3* and *AGO4* | *AGO1*, *AGO3* and *AGO4* |
| **Variation status** | | | *ND* | *de novo* | *de novo* | *de novo* | *de novo* |
| **Current age** (years) | 9.9 | 9.33 | 3.75 | 10.5 | 18 | 1.4 | 13 |
| **Gender** | M:17/F:16 | | | | | | |
| **ID/DD** | 31/31 (100 %) | 3/5 (60 %) | ND | mild | moderate | ND | moderate |
| **Hypotonia** | 13/18 (72 %) | 5/5 (100 %) | yes | yes | yes | yes | yes |
| **Seizures** | 13/28 (46 %) | ND | ND | ND | ND | ND | ND |
| **Motor delay** | 28/30 (93 %) | 4/5 (80 %) | yes | yes | yes | yes | no |
| Walk (months) | ~ 25 | 24 | ND | 21 | 36 | ND | 16 |
| **Speech impairment** | 30/30 (100 %) | 5/5 (100 %) | yes | yes | yes | yes | yes |
| 1st words (months) | ~ 31 | 25 | 18-24 | 24 | 36 | ND | 18 |
| **Autistic behaviour** | 24/30 (80 %) | ND | ND | ND | ND | ND | ND |
| Stereotypies | 11/14 (78.5 %) | ND | ND | ND | ND | ND | ND |
| Hyperactivity | 15/22 (68 %) | 1/1 (100 %) | ND | ND | ND | ND | yes |
| Anxiety | 7/8 (87.5 %) | ND | ND | ND | ND | ND | ND |
| Aggressiveness | 11/14 (78.5 %) | ND | ND | ND | ND | ND | ND |
| **Feeding difficulties** | 10/23 (43.5 %) | 5/5 (100 %) | yes | yes | yes | yes | yes |
| **Sleeping disturbance** | 17/22 (77 %) | ND | ND | ND | ND | ND | ND |
| **Facial findings** | Yes | yes | yes | yes | yes | yes | yes |
| **Other symptoms** | | | astigmatism | buccofacial dyspraxia | bilateral hip dislocation, joint laxity | hypospadias | joint laxiy |
